# Ectomycorrhizal networks propagate carbon deficit and drought stress among trees

**DOI:** 10.64898/2026.05.07.723550

**Authors:** Gerard Sapes, Mary Ellyn DuPre, Alex Goke, Roger Koide, Lorinda Bullington, Anna Sala, Ylva Lekberg

**Affiliations:** Division of Biological Sciences, University of Montana, Missoula, MT 59812, USA; CREAF, Cerdanyola del Valles, 08193 Barcelona, Spain; MPG Ranch, Missoula, MT 59801, USA; Department of Biology, Brigham Young University, Provo, UT, 84602, USA; W.A. Franke College of Forestry & Conservation, University of Montana, Missoula, MT 59812, USA

## Abstract

How plants allocate carbon determines their productivity, responses to stress, and interactions with other organisms. A substantial amount of plant carbon is stored as non-structural carbohydrates (NSC), which sustain turgor via osmoregulation and fuel metabolism when carbon is limited. NSC also support root-colonizing mycorrhizal fungi, thus we hypothesized that under carbon-limiting conditions such as drought, a trade-off between feeding mycorrhizal fungi and maintaining turgor may arise. We reduced carbon allocation to ectomycorrhizal (EcM) networks by girdling *Pinus ponderosa* trees exposed to drought or ambient conditions and manipulated putative fungal connections between trees by trenching. We show that, in droughted plots, trees putatively connected to girdled trees by EcM networks had 33 % less needle NSC and >10% less turgor than those connected to ungirdled trees. Trees disconnected from the mycorrhizal network by trenching had increased NSC likely from the increased water availability with girdling, but these gains were offset in the presence of networks. Our results demonstrate that the increased carbon demand by EcM fungi in response to reduced carbon inputs from some trees can deplete NSC in neighboring trees via shared mycorrhizal networks. At least in the short term, allocation trade-offs under carbon-limiting conditions may expose networked trees to carbon deficits. This may increase vulnerability to drought, which may be particularly acute given shifts in climate.

## Introduction

How plants allocate carbon is a central determinant of their performance, their responses to stress, and their interactions with symbiotic organisms^1^. In addition to carbon required for growth, energy, reproduction, and defense, plants store carbon as non-structural carbohydrates (NSC)^2^. NSC are thought to serve both as a carbon buffer during periods when carbon demand exceeds supply^3^ and as a source of osmolytes to constantly maintain turgor^4^. Plants also transfer carbon belowground for root growth and to sustain obligate symbionts and other soil biota^5–7^. Drought, high temperature, defoliation, fire, and other stressors can reduce carbon supply relative to demand and prompt allocation tradeoffs among sinks. While tradeoffs involving growth have received substantial attention^8–10^, we know substantially less about priorities of other sinks. Here we focus on potential trade-offs between two of these sinks: carbon allocation to NSC storage and to symbiotic ectomycorrhizal (EcM) fungi, and the impact that such trade-off might have on carbon-dependent plant water relations such as osmoregulation. An increased knowledge in this area is critical for modelling forest responses to stress due to climate change.

Plants store approximately 10% of their biomass as NSC in the form of starch and soluble sugars, and this storage is rarely reduced below ∼40% of the seasonal maximum^2^. While these abundant NSC pools could indicate that mature trees are not carbon limited^11–13^, they may also suggest additional vital functions of NSC^14,15^. For example, it is well known that plants use NSC for the synthesis of compounds^16–18^ that are critical for metabolic functions, osmoregulation, and energy for active transport^2^. What is less known, however, is how much of the stored NSC are required for turgor maintenance, including its energetic costs. Turgor maintenance is thought to be very costly^19^, although direct empirical evidence is rare^4^. Sapes et al. (2021)^4^ showed that experimental depletion of NSC below ∼40% of reference control compromised cell turgor and osmotic capacity in well-watered seedlings in the greenhouse. Similar effects were recently observed in adult trees under natural conditions (Sala et al., *in prep*), which highlights the intricate link between NSC storage and plant water relations. Substantial carbon costs associated with turgor maintenance could explain why the size of stored NSC is under selective pressure^10,20^ and the high estimated cost of osmoregulation in models^19^. Quantifying the carbon cost associated with drought responses, and assessing trade-offs with other sinks is pressing to better gauge the ability of plants to respond to stress.

In temperate and boreal coniferous ecosystems, virtually all trees are colonized by ectomycorrhizal (EcM) fungi, and this symbiosis can increase access to water and nutrients^21–25^. However, these benefits come at a cost to the plant as EcM fungi receive an average of 13% (and some up to 50%) of assimilated carbon^6,26,27^. Fungi can colonize multiple trees, connecting them in complex networks^24^, while acting as strong sinks^28–31^. This opens the possibility of unequal carbon contribution from plants to the EcM network based on tree phenology, size, and health status^32^. Indeed, a greenhouse experiment with seedlings showed that healthy neighbors connected to carbon-stressed neighbors became NSC depleted^4^, presumably because the carbon demand by EcM fungi on the healthy seedlings increased due to reduced contribution by the carbon-stressed neighbor. This coincided with a decreased ability by the healthy seedling to osmoregulate and maintain turgor, indicative of a carbon allocation trade-off^4^. Thus, stress was propagated via the EcM network but, importantly, did not involve or require host-to-host carbon transfer, which currently has little support in field studies ^33^. Individual trees in forests may be lost to windfall, pests, or selected logging. While this may reduce competitive pressure on neighbors^34^, no study has examined the alternative outcome that carbon demand from EcM fungi increases—at least temporarily—which could reduce the ability to tolerate drought of surviving trees.

Under field conditions, we examined whether EcM networks can propagate NSC depletion among neighbor trees and contribute to drought vulnerability in the widely distributed ponderosa pine (*Pinus ponderosa*). To do so, we established plots consisting of groups of three ca. 18 year-old ponderosa pine trees that were sufficiently close in proximity to be connected via EcM fungi (Fig. 1) and where half of the plots experienced artificial drought while the rest experienced ambient precipitation. We girdled one tree to eliminate NSC supply to EcM fungi, measured responses in a second tree, and asked if responses differed from those in the third tree where putative EcM network connections to the girdled tree had been ruptured via trenching. These responses were then compared to control plots where no trees were girdled (Fig. 1). Additionally, because EcM networks are nearly impossible to document directly in the field^33^, in ambient plots only we performed an add-on experiment with seedlings planted next to girdled trees in static or rotated cores that either allowed or severed connections with EcM fungi outside the cores^35^. We hypothesize that the reduced carbon supply to EcM fungi associated with girdling would increase the EcM fungal carbon demand and reduce stored NSC in neighbor trees and seedlings that were still connected to the EcM network. This, in turn, would lead to lower ability to osmoregulate and maintain turgor, especially in the drought treatment. It is well documented that EcM fungi can improve plant performance during drought via a plethora of mechanisms^36^. Whether potential negative effects also exist under field conditions and can be propagated via mycorrhizal networks is not known. Such knowledge is critical given the dominance of EcM fungi in temperate and boreal forest and the increased drought and heat stress predicted under climate change.

**Figure 1.**
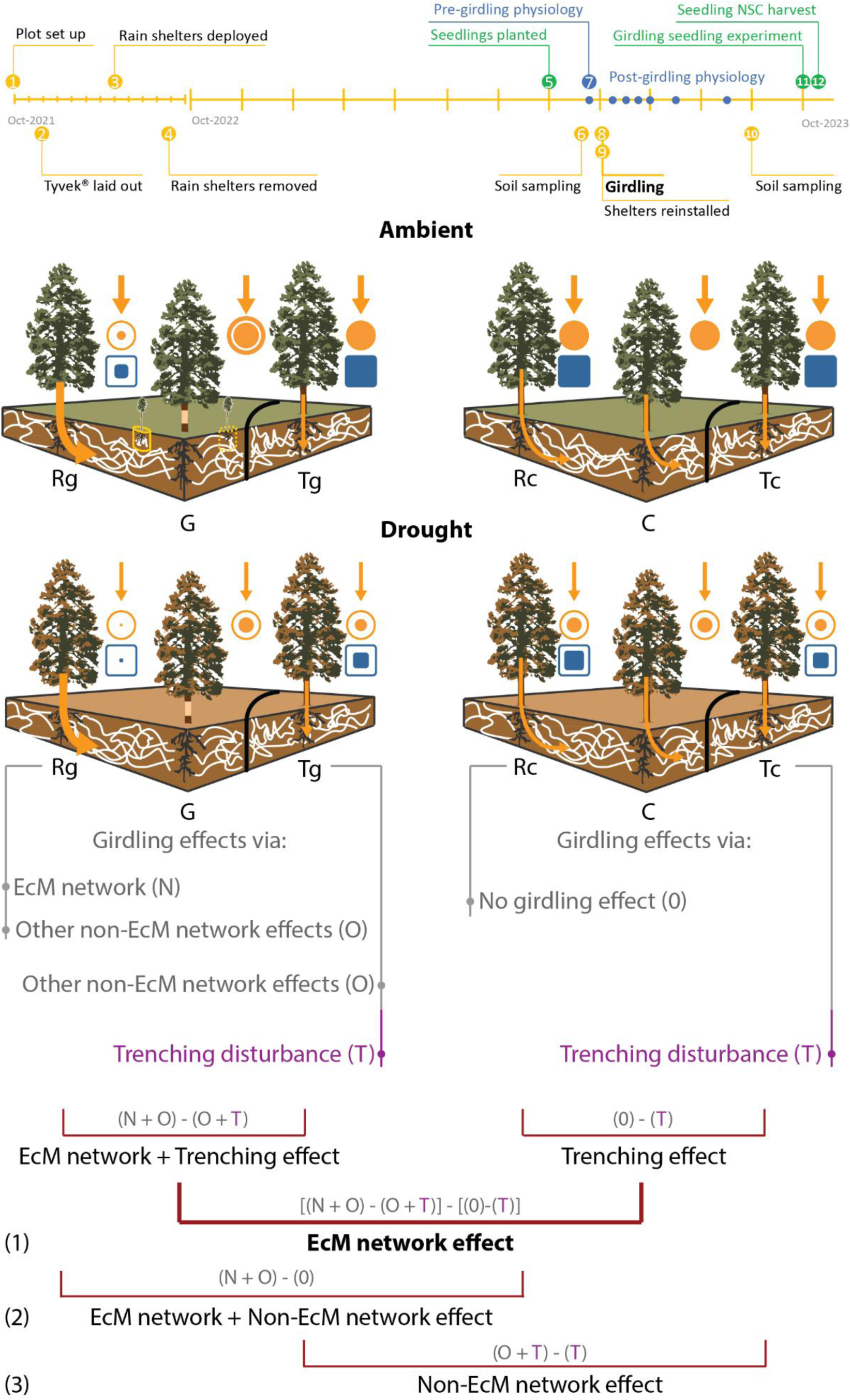
Experimental design and hypotheses. Top: Timeline of events for the experiment. Each tick mark corresponds to a month. The timeline for the 2023 experiment is zoomed in for readability purposes. Bottom: Green and brown plots represent ambient and drought treatments, respectively. Middle trees with striped bark and whole bark represent girdled (G) and control (C) trees, respectively. Left-side trees represent response trees potentially connected to girdled (Rg) or to control (Rc) trees via network-forming ectomycorrhizal fungi (white). Right-side trees represent trenched trees next to girdled (Tg) or control (Tc) trees disconnected from the ectomycorrhizal network by trenches (thick black line). The plots were also surrounded by a trench (thin black line). We hypothesize that girdling decreases carbohydrate allocation belowground to EcM fungi causing demand to increase from R trees connected to the same mycorrhizal network. This results in decreased NSC storage in R trees (partially filled orange circle) with negative consequences on turgor maintenance (partially filled blue circle). If these effects are mediated by EcM networks, they should be absent in T trees as they are not connected to the EcM network. We also hypothesize that effects are exacerbated under prolonged drought due to reduced carbon assimilation. Effects were quantified using differences between T and R trees (small thin red boxes) in girdled and control plots and by comparing those differences in ambient vs. drought plots (thick red box, see below for details). Text under plots show ways through which girdling (gray) and trenching (purple) can influence R and T trees. Rg trees can be affected by girdling through EcM network (EcM network effects) and soil pathways (Non-EcM network effects). Tg trees are isolated from the network and can only be affected by soil pathways. Rc and Tc trees cannot be affected by girdling through any of these pathways because their neighbor was not girdled. Any response in T trees could however be due to unexpected trenching effects that must be accounted for. Equations 1, 2, and 3 describe how EcM and Non-EcM network effects can be estimated by strategically subtracting these effects among tree types and treatments.

## Results

### Modest treatment effects on EcM fungi

In May 2023, prior to girdling, fungal biomass was greatest at 0-10 cm and declined drastically with depth in control plots (F _(3, 14)_ = 25.9, *p* < 0.001, Fig 2a), indicating that our trenching depth was sufficient to disrupt most EcM networks. Also in May 2023, fungal biomass was 57.0% ± 16.3% lower in drought plots than ambient plots, but only in the 0-10 cm depth (t= 2.3, *p* = 0.04), indicative of a legacy of the experimental drought that occurred in 2022. These suppressive effects of drought had disappeared by September (F _(1, 16)_ = 1.3, *p* = 0.2) and girdling did not affect fungal biomass (F _(1, 16)_ = 0.8, *p* = 0.3, Fig 2b). Similarly, we observed no difference in EcM fungal relative sequence abundance across girdling (F _(1, 32)_ = 2.8, *p*= 0.1) or drought (F _(1, 32)_ = 1.6, *p*= 0.2), which averaged ∼30%.

**Figure 2.**
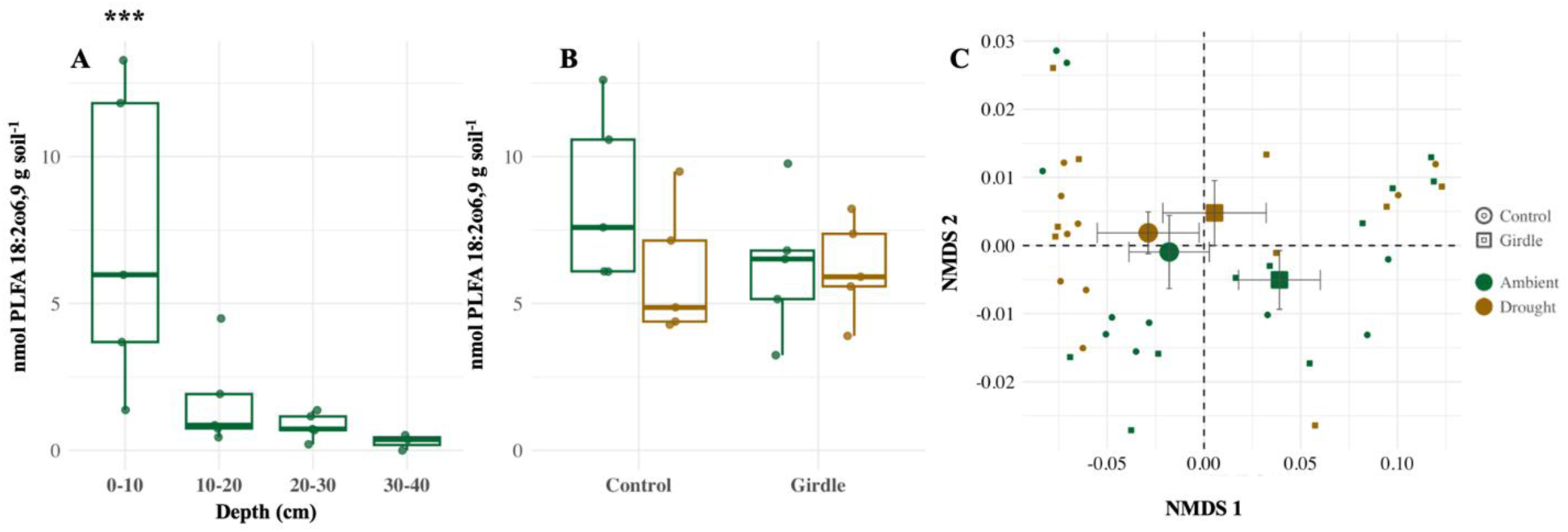
Fungal biomass declined with depth, but EcM fungal communities were resistant to drought and girdling. Fungal biomass measured across 10 cm depths in control (non-girdled) plots in May 2023 (a), and in control and girdled plots in September 2023 (b). NMDS of EcM fungal community composition in soil collected in girdled or control plots experiencing drought or ambient conditions (c). In (a) and (b), Asterisks indicate significant (p<0.05) differences between depth levels, center lines of boxplots indicate medians, boxes represent interquartile ranges, whiskers extend to the most extreme values within 1.5 × the interquartile range, and points show individual observations. In (c), large symbols denote treatment mean NMDS scores, error bars indicate ± standard error, and small points represent individual observations. Significance levels: · = p < 0.1, * = p < 0.05, ** = p < 0.01, *** = p < 0.001

EcM fungal community composition did not differ among control plots (F _(1, 15)_ = 0.6, R^2^=0.04, *p* = 0.6) or between C and Rc trees in September, 2023 (F _(1, 15)_ = 0.9, R^2^=0.02, *p* = 0.9, Fig. S4). This indicates no strong spatial structuring among and within plots and thus a greater likelihood of networks to form among neighboring trees. Also in the control plots, there was no effect of trenching because we found no difference in communities in soil collected near Tc and Rc trees (F _(1, 15)_ = 0.4, R^2^=0.02, *p* = 0.7).

Dominant EcM fungal genera in control plots included *Wilcoxina* (38% of EcM fungal sequences), *Inocybe* (18%) and *Rhizopogon* (17%). *Wilcoxina* and *Inocybe* are characterized as short-distance exploration types that do not form rhizomorphs, whereas *Rhizopogon* is a long-distance exploration type that form rhizomorphs^37^. Less abundant rhizomorph forming genera (<5% of EcM fungal sequences) included long-distance *Suillus* and medium-distance *Amphinema*, *Cortinarius*, *Lactarius*, *Lyophyllum*, *Piloderma*, *Polyozellus*, *Pseudotomentella*, *Thelephora*, *Tricholoma*, and *Tomentella*.

Drought did not affect the EcM fungal community composition (F _(1, 32)_ = 1.07, R^2^ = 0.03, *p* = 0.3) and effects of girdling were weak (F _(1, 32)_ = 3.1, R^2^=0.08, *p* = 0.05, Fig. 2C). One ASV in the genus *Inocybe* (a short-distance exploration type) was relatively more abundant in girdled than control plots (lfc = 3.16, *q* = 0.002). Richness of EcM fungi was marginally lower in response to drought (-3.3± 2.08 ASVs; F _(1, 33)_ = 3.6, *p* = 0.1) whereas girdling had no effect (F _(1, 33)_ = 0.1, *p* = 0.7).

### Girdled trees became NSC depleted

Girdling caused an immediate drop in phloem soluble sugars below the girdle wound of the girdled trees in both ambient and drought plots (Fig. S5A-B, Table S1A & S1D). After the immediate drop from the initial 7.5 %, soluble sugars tended to stabilize close to a minimum threshold of 2.5%. Starch and total NSC pools also declined (Fig. S5C-F, Table S1B-C & S1E-F) over the course of two and a half months, but there was no immediate depletion as observed for soluble sugars. Depletion rates below the girdling wound did not differ between ambient and drought plots (Table S2).

### Girdling also impaired neighbor trees

The difference (R_G_-T_G_) - (R_C_-T_C_), which measures EcM network effects on neighbor NSC and water relations after girdling while accounting for the trenching disturbance (Fig. 1; see *Statistical Analyses*), was significant in drought plots only. Specifically, once accounting for potential side-effects (if any) of trenching in girdled and control plots, soluble sugar concentrations were lower in neighboring response trees in girdled plots than those in control plots at 14 d (difference of -0.814 ± 0.322 % free sugars) and 28 d (decrease of 0.795 ± 0.348 % free sugars; Fig. 3B, Table S4D) after girdling. This difference represents a ∼33% reduction from control drought plots where the concentration of soluble sugar in needles was ∼2.5%. Trees in ambient and drought girdled plots did not differ in starch or total carbohydrates relative to control plots. Once accounting for trenching effects in girdled and control plots, pressure potential was also lower in response trees in girdled plots relative to response trees in control plots 6 d after girdling a neighbor (difference of 0.250 ± 0.105 MPa) and trended lower at 28 d (difference of 0.174 ± 0.078 MPa, Fig. 3D, Tables S3D). This represents a reduction in turgor of ∼12% relative to the control on day 6. We did not detect statistically significant differences in osmotic potential. However, we observed a trend of increased osmotic potential on the days we detected significantly lower turgor pressure. Water potential was marginally lower at 6 d in response trees from girdled plots relative to control plots, once accounting for trenching effects. Overall, girdling a tree had negative effects on neighboring, putatively connected trees, but only when trees experienced drought.

**Figure 3.**
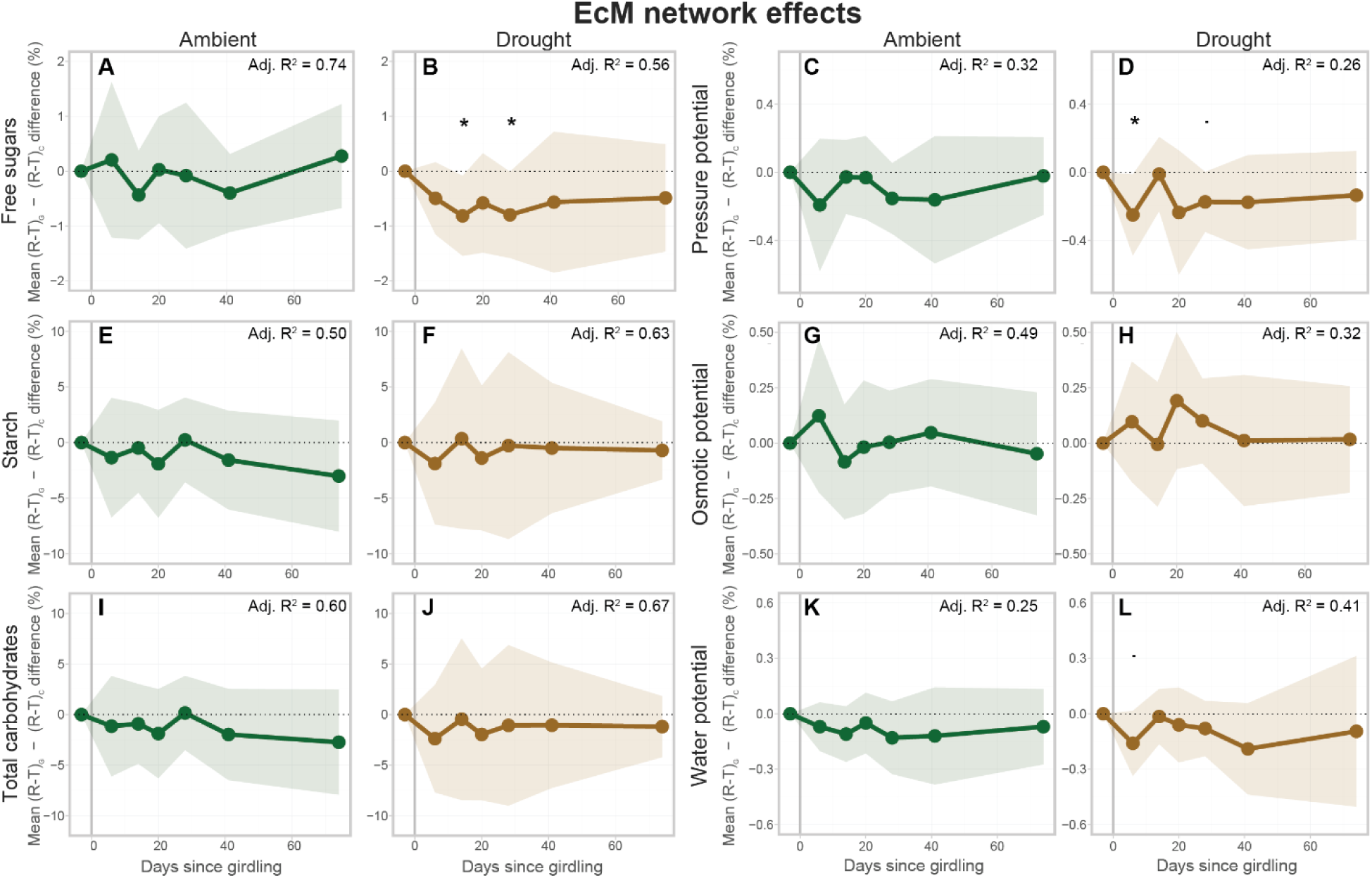
Putative EcM networks spread carbohydrate depletion and impact water relations under drought. Panels show DiD estimates of treatment effects in non-structural carbohydrates (columns 1 and 2) and water relations (columns 3 and 4) under ambient (green) and drought (brown) conditions. Vertical line indicates the start of treatment. The DiD estimate is the difference in differences between response and trenched trees between girdled and control plots. Shaded areas represent 95% mean confidence intervals. Significance levels: · = p < 0.1, * = p < 0.05.

### Girdling improved trenched trees’ status

The DiD models in Fig. 3 assume that the trenching effect on T trees is the same in girdled and control plots. However, girdled trees may take up less water and nutrients^38^, which could benefit neighbor trees (both Tg and Rg trees) via competitive release. If so, any negative responses in Rg trees could be offset by the increase in water availability due to girdling-induced competitive release. Note that this effect is not a trenching treatment artifact but instead represents a non-EcM network effect resulting from girdling. To quantify this effect, we first compared Tg and Tc directly (Tg – Tc). Six days after girdling, Tg had higher pre-dawn water potential than Tc in drought plots (Figs. 4, S8). This indicates that girdling initially increased water availability in neighbor trenched trees (Fig. 4A). This was followed by an increase in soluble sugars in Tg relative to Tc that was sustained over time (Fig. 4C, Fig. S6, Table S6D). Specifically, we detected an increase in absolute sugar concentration in Tg relative to Tc of 0.551 ± 0.299 %, 0.545 ± 0.244 %, 0.863 ± 0.256 %, and 0.869 ± 0.305 % of dry mass, 6d, 14d, 28d, and 74 d after girdling, respectively. These values correspond to a relative increase in soluble sugar concentrations of 24.20 %, 22.38 %, 50.01 %, and 51.68 % relative to Tc at 6d, 14d, 28d, and 74d after girdling, respectively. Predawn water potential was also 0.140 ± 0.073 MPa higher in Tg than Tc at 41 days after girdling in ambient plots (Fig. S6; Table S5C & F). In sum, trenched trees in girdled plots had access to more water and contained more soluble sugars compared to trenched trees in control plots, but this effect was almost exclusively observed under drought.

**Figure 4.**
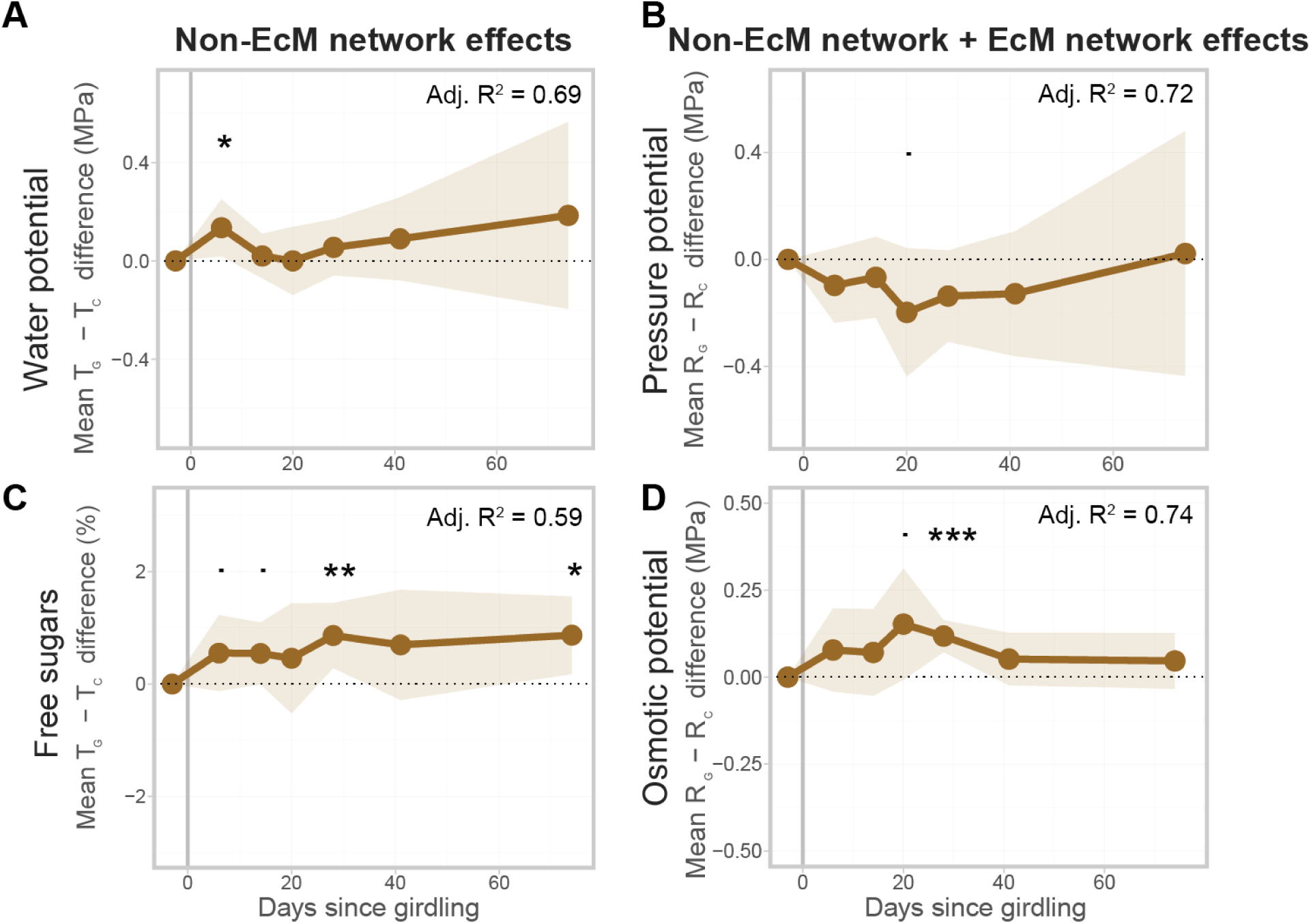
Under drought, girdling improved access to water and photosynthate production in trees isolated from the EcM network and impaired water relations in neighboring response trees. Panels show DiD estimates of treatment effects in water potential (A), pressure potential (B), free sugars (C), and osmotic potential (D) in trenched (left) and response (right) trees. Panels A and C represent non-EcM network effects while panels B and D represent both non-EcM network effects and EcM network effects. Vertical line indicates the start of treatment. The DiD estimate is the difference between either trenched or response trees in girdled and control plots. Shaded areas represent 95% mean confidence intervals. Significance levels: · = p < 0.1,* = p < 0.05, ** = p < 0.01, *** = p < 0.001.

We then calculated differences between Rg and Rc (Rg – Rc), which do not differentiate between EcM and non-EcM network effects of girdling. If there are no EcM network effects, we expect the same differences in R as in T trees. However, contrary to what we observed in T trees, Rg tended to have *lower* pressure potential 20 d after girdling than Rc in drought plots (-0.198 ± 0.106 MPa, Fig. 4B, Fig. S7, Table S7D). Also, osmotic potential was up to 0.152 ± 0.071 MPa higher (less negative) in Rg than Rc with marginally significant differences at 20 d and highly significant differences at 28 d (Fig. 4D, Fig. S7, Table S5E). Thus, while trenched trees benefited from a girdled neighbor under drought, response trees with putative EcM network connections with the girdled neighbor appeared to be harmed.

### EcM networks suppressed seedling NSC

Seedling root samples were colonized by an average of 4.55 ± 0.44 EcM fungal ASVs, with communities differing among ambient plots (F _(1, 19)_ = 2.1, R^2^ = 0.65, *p* = 0.04) but not between static or rotated cores (F _(1, 19)_ = 0.5, R^2^ = 0.28, *p* = 0.6), indicating that core treatment differences are unlikely to be a major driver of variation in downstream NSC and δ¹⁵N analyses. Like soil samples, ASVs in the genera *Wilcoxinia* (33% of sequences), *Rhizopogon* (25%), and *Thelephora* (14%) were prominent.

δ^15^N was higher in the static than rotated cores (F_(1,18)_ = 5.3, p = 0.03), which did not differ from natural abundance. This suggests that EcM fungal hyphae outside the core had grown into the core and colonized seedlings whereas those in the rotated cores did not (Fig. 5b).

**Figure 5.**
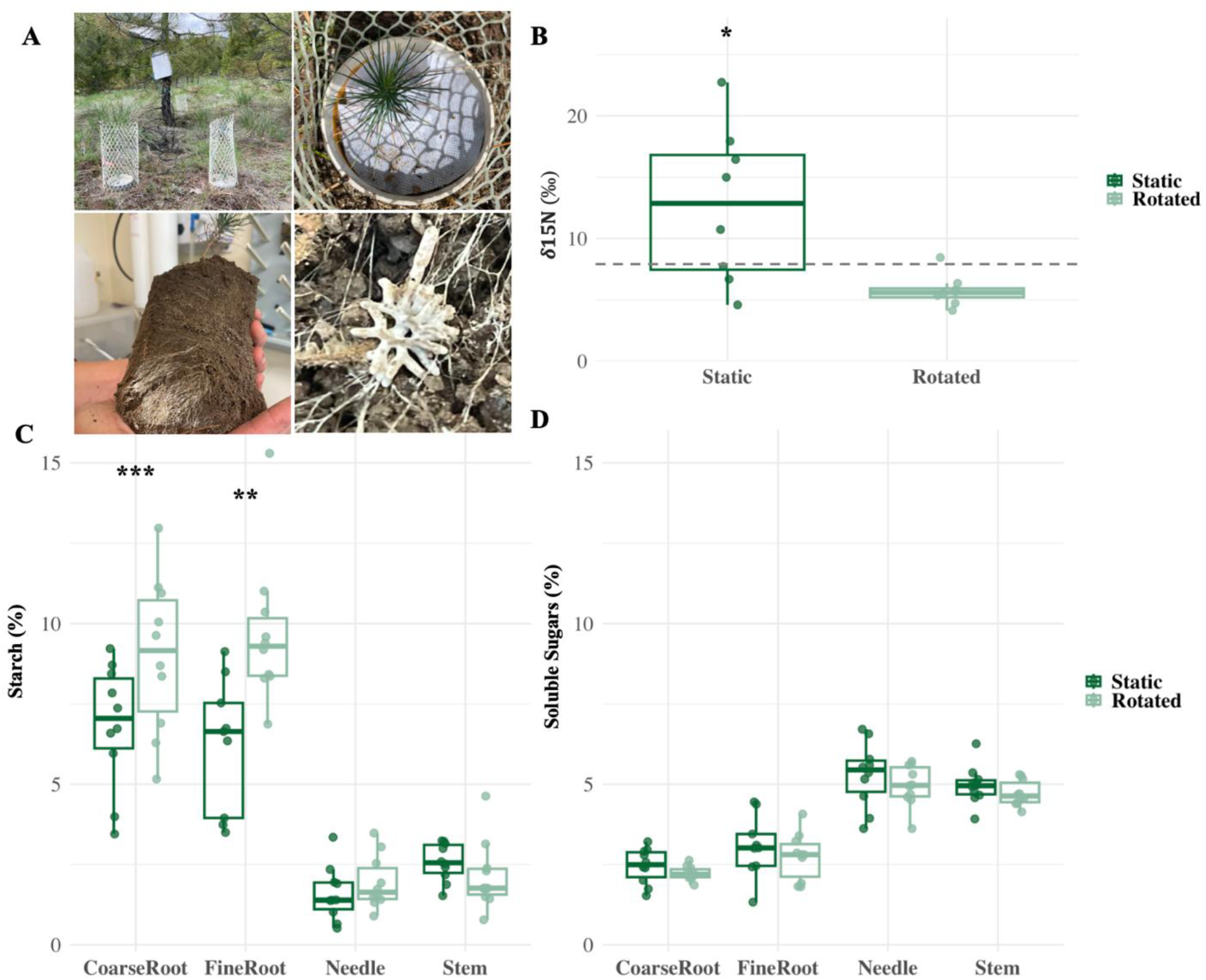
Planted seedlings were colonized by EcM from outside cores and seedlings in static cores had lower starch concentrations. Seedling experiment where one out of the two cores was rotated after five months following visual colonization by surrounding EcM fungi, including rhizomorph-forming EcM (A). δ_15_N values at harvest in EcM root tips associated with seedlings grown in static or rotated cores (B), and comparisons of starch (C) and soluble sugars (D) in tissue types. Asterisks indicate significant (p<0.05) differences between static and rotated cores, n=10. Dashed line in panel B indicates natural abundance of δ_15_N. Significance levels: · = p < 0.1, * = p < 0.05, ** = p < 0.01, *** = p < 0.001

NSC in needle and stem tissue did not differ between static and rotated cores, but coarse and fine roots in the static cores had fewer total NSC compared to roots in the rotated cores (F_Core x Tissue type (3, 71)_ = 5.1, *p* = 0.003). This decline in NSC in the static cores was driven by starch (F_Core x Tissue type (3, 71)_ = 5.6, *p* = 0.002; Fig 5D), where starch was 24.2 ± 8.2 % lower in fine roots (t = -3.5, *p* = 0.004) and 35.6 ± 7.8 % lower in coarse roots (t= -2.2, *p* < 0.001) growing in static compared to rotated cores. Soluble sugars differed by tissue type (F_Tissue type (3, 71)_ = 81.9, *p* < 0.001) but did not differ between static and rotated cores (Fig. 5D).

Thus, similar to adult trees next to girdled neighbors, seedlings had more NSC when putative EcM network connections were severed.

## Discussion

We show here that reducing carbon allocation belowground from one tree caused reductions in NSC, osmoregulation capacity, and turgor in a neighbor tree in droughted plots. This effect was not observed when neighbors were separated from girdled trees by a >30 cm deep trench severing putative EcM connections. Also, seedlings growing next to girdled trees with intact connections to the EcM network had lower NSC concentrations in their roots than seedlings where EcM connections had been ruptured. While EcM networks are notoriously difficult to research *in-situ*, we argue that the most parsimonious explanation to our findings is that the reduced carbon transfer to EcM fungi by girdled trees increased carbon demand from neighboring trees connected to the EcM network, at least temporarily. This reduced the capacity to lower the osmotic potential in neighbor trees, indicative of a carbon allocation trade-off that manifested when trees were drought stressed. The trade-off only arose under drought, when carbon limitation was likely higher. Context dependency is a classic feature of resource allocation trade-offs. Below, we outline these mechanisms in more detail, explore alternative explanations, debate whether this trade-off challenges the altruistic frameworks on the relationship between trees and mycorrhizal fungi^39^, and outline implications for forest management and resilience in light of ongoing climate change.

### Alternative explanations to EcM networks

Because continuous mycelial connections among trees are near impossible to track in the field^33^, we must contemplate alternative explanations before attributing observed effects solely to EcM networks. Our results cannot be due to drought-mediated changes in source-sink dynamics because these effects would affect all trees within drought plots, not only response trees. Our findings in Rg trees can also not be due to water limitation resulting from greater uptake by Tg trees because this should lead to osmotic adjustment through increased sugar concentrations and lower osmotic potential. Instead, we observed the opposite pattern which was also seen in greenhouse conditions^4^. However, we can think of two additional mechanisms that could generate stronger carbon sinks near response trees next to girdled trees only. First, carbon depletion in roots of girdled trees could trigger NSC transfer from neighbor trees via root grafts^40^. Root grafts have been observed in closely related conifer species, but often in older trees growing at higher densities than trees in our plots^41^. We found no reports of root grafts in ponderosa pine in the literature, and no root grafts were found when we excavated five pairs of ponderosa pine trees outside our plots (Appendix S1). Also, no movement of dye to neighbor trees was noted after acid fuchsin injections^25^ into stumps (Appendix S1). Based on this, movement of NSC among trees through root grafts seems unlikely. Second, some root exudation is source-sink driven and soil biota can be strong sinks^42^. It is possible that girdling reduced exudation and this increased the demand by rhizosphere organisms from neighbor trees if root systems overlapped. This may be exacerbated by drought as it can increase exudation of signal molecules that attract mycorrhizal fungi^43^. Although we cannot unequivocally exclude this possibility, NSC concentrations also decreased in seedlings where EcM network connections—and not root associations—were manipulated. Also, carbon allocation to EcM fungi can be five times greater than carbon lost to root exudation^44^. Based on this, and the similarity of EcM fungal communities in soil next to neighboring trees—including rhizomorph forming taxa known to form EcM networks^45^—we argue that an increased carbon demand by EcM networks is the most likely explanation for our results. This indicates that patterns previously observed in the greenhouse^4^ where EcM networks propagated stress from one seedling to another also happen in both seedling and adult wild trees in the field.

### Context-dependent girdling responses

In contrast to the idea that stored NSC may reflect surplus carbon^1^, our results suggest a context-dependent carbon allocation tradeoff between maintaining osmoregulation and ectomycorrhizal symbionts in response trees experiencing drought. Specifically, low NSC concentrations were associated with impaired water relations under drought, but only in neighbor trees not separated from girdled neighbors by a trench. Trenched trees, on the other hand, had 0.14 MPa higher water potential which likely resulted in the observed **36%** increase in free sugar concentrations relative to trenched trees in control drought plots. This may be due to increased water availability in girdled plots, because girdling can cause feedback inhibition of photosynthesis and stomatal closure^46^. That raises the question; if water was more available in girdled drought plots, why were response trees in those plots more drought stressed (i.e., lower turgor) than response trees growing next to ungirdled, drought neighbors?

The direct comparisons of response trees in girdle and control plots represent the combined impacts of putative EcM and non-EcM network effects (Eq. 2 in Fig. 1, Fig. 4B&D) where any gain in carbohydrates or water relations from increased water availability due to girdling can be offset by losses due to increased carbon demand by EcM fungi. In our study, the negative effects by EcM fungi surpassed benefits from competitive release, leading to an overall ca. **8%** increase in osmotic potential and a ca.

**11%** decrease in pressure potential in response trees in girdled plots relative to control plots (Fig. 4 B, D). Isolating the EcM network effect (Eq. 1 in Fig. 1, Fig. 5) yielded an estimated NSC depletion attributable to EcM networks of 33% (calculated as the estimated 0.814% sugar concentration depleted divided by the ca. 2.5% existing in control drought plots at day 14 after girdling). This depletion value is similar to that observed in response seedlings in Sapes et al. (2021)^4^ and right below the seasonal NSC depletion levels relative to seasonal maximums observed globally^2^. The 12% impairment in pressure potential attributable to EcM networks is also similar to that of Sapes et al. (2021)^4^. Because this is predawn turgor when drought stress is at its daily minimum, these estimates of physiological stress are likely conservative and may worsen as water potential declines throughout the day. Likewise, while negative effects were only statistically significant during some sampling events following girdling, free sugars and pressure potential showed sustained trends of decline until the end of the summer season when we stopped measuring. This lack of significance may at least partly be due to low statistical power resulting from compounding errors from four mean estimates (Rg, Tg, Rc, Tc) which led to wide confidence intervals, limited number of replicates, and intrinsic noise typical of field experiments. Yet, we still detected a treatment signal. Thus, reduced carbon allocation belowground from neighbor trees in natural stands could negatively impact functions such as carbon assimilation, growth, reproduction, defense, and drought tolerance that depend on NSC status and turgor^47^, especially if we consider the cumulative effect of being several weeks under depressed turgor as observed in our trees (Fig. 3D).

While trees and seedlings showed similar reductions in NSC, they differed in the type of NSC that was depleted and whether this affected water retention capacity. Soluble sugars were depleted in trees whereas starch was depleted in seedlings. Our previous research with seedlings in the greenhouse also showed starch depletion and stronger relationships between osmoregulation capacity and starch than sugars^4,48^. We also observed starch depletion and impacts on osmotic capacity in a recent field experiment where shading caused severe carbon limitation. Thus, starch degradation may only be mobilized when EcM fungal carbon demand is high enough to risk compromising sugar concentrations^44^. This might occur more often in seedlings than trees where carbon allocation trade-offs may manifest sooner, possibly due to more frequent carbon limitation due to demand from growth and reduced NSC pools. Contrary to trees in our study, the lower NSC concentrations in seedlings connected to the EcM network did not affect water retention capacity. This is likely because seedlings were not drought stressed as they were grown in ambient plots and watered when needed.

### Biological relevance and paths forward

The wood-wide-web publication by Simard et al. 1997^39^ highlighted the interconnectedness in forests, with EcM networks potentially allowing sharing of resources and signals among trees^49^, kin-recognition^50^, and improved seedling establishment^51^. While these findings have been questioned recently, mycorrhizal networks clearly have the potential to influence plant-plant interactions in both positive and negative ways^52^. Here, we showed that EcM networks can promote negative plant-plant interactions at times of stress in unexpected ways by affecting host carbon status without requiring plant-to-plant transfer, which is controversial^33,53^. While we reduced carbon flux into EcM networks by girdling trees, similar effects could result from partial stand mortality from spotty fires, insect and fungal outbreaks, thinning, and droughts. A reduced stand density may benefit remaining trees as more resources are available, but increased carbon demand from EcM networks could at least temporarily counteract this. For example, thinned ponderosa pine trees fared better during drought than non-thinned stands using aerial thermal imaging and hyperspectral data^54^, but ponderosa pine in our study became more stressed. It is paramount to understand when, where, and how EcM networks might negatively impact trees through their interactions. Doing so requires studying them under shifting environmental conditions given their highly dynamic nature.

Our study also invokes many questions about the mycorrhizal fungi themselves. First, do EcM networks persist for long periods or do they rebuild every year? While longevity of fungi is notoriously difficult to ascertain, mesh bag studies have indicated a turnover of multiple times per year^55,56^, although this may reflect fine mycelium rather than the whole network. Size and longevity of mycelia likely range widely among EcM fungal taxa^57^, but rebuilding from scratch every year seems unlikely based on comparisons of mycelial production relative to standing biomass^56^. Thus, while EcM fungal biomass likely peaks in the fall at our site and elsewhere^56,58,59^, we documented patterns consistent with EcM network connections in early summer indicating a persistence of functional networks across seasons. We also documented high resilience in these fungal communities, because differences in fungal biomass among treatments had disappeared in the fall, and drought did not affect the composition of communities, which contrasts with some previous work^60–62^.

Our findings are tantalizing, but more research is needed to establish long-term impacts of EcM networks on tree performance and vulnerability to drought, and to assess their generality as not all trees or species within a network may respond the same^63^. For example, if some trees supply more carbon to the EcM network than others, removal of those trees may cause larger demand when they die, making it more challenging for remaining trees to offset deficits. Similarly, EcM networks composed of smaller number of trees may be more vulnerable to partial stand mortality given that the carbon deficit left by a dying tree would be distributed among fewer trees. The interdependencies between plant carbon, osmoregulation, and symbiont maintenance evidenced here provide an interesting opportunity to incorporate interactions between trees and belowground symbionts into carbon models through the already known mechanistic relationships between NSC and water relations. Also, more information about mycelial growth and turnover would allow inclusion of mycorrhizal fungi into ecosystem models^64^.

## Conclusions

Overall, we have shown here that EcM fungi can make trees in the field more vulnerable to drought, at least temporarily. However, EcM fungi can increase drought tolerance through various mechanisms^36,45^, and it is also possible that compensating for temporary reductions in carbon allocation from dying trees by other hosts within the network may increase long-term forest resilience by maintaining EcM networks and their benefits through periods of stress. Understanding under what conditions EcM fungi promote drought tolerance vs. exacerbate stress is imperative, and our findings highlight some of the complexity underlying these different outcomes. Understanding these dynamics is increasingly pressing given ongoing climate change, which is predicted to increase the frequency and duration of drought and cause forest dieback^65^, including in the >56 million ha of ponderosa pine forests across the western USA that currently experience a megadrought^54^. Mycorrhizal fungi, and root-associated fungi in general, should thus be considered to better understand how trees may respond to drought as plant adaptation to drought may depend on mycorrhizas^66^.

## Materials and Methods

### Experimental design, site, and plot set up

The experiment was set up at Woodchuck Gulch in the Blackfoot Valley of western Montana (46°58’06"N, 113°25’39"W) (Fig. S1A). This site is a dry, mostly flat, open ponderosa pine forest at 1200 m of elevation with abundant natural regeneration after restoration harvesting in the early 2000s. Mean annual temperature at Seeley Lake (Montana, USA), ca. 20 km from the site, is 5.1 °C, with an average maximum temperature of 28.1 °C in July, and an average minimum of -11.2 °C in January. Mean annual precipitation is 388 mm, with the wettest months in May and June. The understory is dominated by grasses (mainly *Festuca campestris*), forbs, and to a lesser degree the ericaceous shrub, *Arctostaphylos uva-ursi*. The site is open to cattle grazing during the summer although an electric fence around plots kept cattle off during the experiment. A weather station assembled with a Campbell CR1000 datalogger, relative humidity and air temperature (Campbell HMP45C), global radiation (LI-COR LI-200), precipitation (Texas Electronics TR-525), and wind velocity (Met One 034E) sensors was installed in a clearing.

The experiment consisted of a 2 x 2 factorial design with water (ambient and drought) and girdling (girdled and control) as main factors with 5 replicates each for a total of 20 plots (Fig. 1). Each replicate plot consisted of three trees where the middle tree was either girdled (G) to prevent carbohydrate transfer to belowground, or not (Control, C). Physiological responses were measured in a response tree (R) on one side of the middle tree (G or C), and in the trenched tree (T) on the other side. The T tree was isolated from the putative EcM network by a narrow 20-30 cm trench dug one week prior to girdling at equal distance from the middle (either G or C tree) and T tree. This trench was refreshed weekly to prevent re-establishment of network connectivity by new fungal growth.

To prepare the plots, we identified 20 clusters of ca. 18-year-old ponderosa pine trees that were at least 6-10 m away from older and taller, mature trees in October 2021. Each cluster had a minimum of three trees at a distance from each other between 0.6 and 4.4 m to maximize the chances of a shared EcM network (Fig. S1B-C). Extra seedlings and saplings were cut leaving only the three target trees. On average, plots were 44 m^2^ (25 - 80 m^2^). We used an iterative randomization process to assign treatments to each plot while minimizing differences in tree size and distances among the three trees within a plot.

ANOVA tests found no significant differences in tree height, diameter, and distance among plots assigned to the four treatments (not shown). We trenched the perimeter of each plot to a depth of 30-45 cm to sever EcM fungal connections to trees outside the plots and to eliminate surface horizontal water flow into plots. Estimates of fungal biomass over soil depth (see below under fungal analyses) confirmed that fungi preferentially occur in the top 10 cm of the soil at this site (Fig. 2; see *Results*), which agrees with previous findings that EcM fungi predominately occupy the top 15 cm of soil^67,68^.

The drought treatment was initiated in December 2021 by placing Tyvek® fabric on the assigned drought plots (total n = 10) to prevent snow percolation (Fig. S1B). Tyvek® is white and impermeable to water but not to gases, thus minimizing soil heating and allowing efflux from soil respiration during snow-free periods. The fabric extended to the outside of the perimeter of the plot to minimize water infiltration and increase the severity of drought. We removed the Tyvek® once the snow had melted and before air temperature increased on May 9^th^. Rain shelters were deployed on May 10^th^ 2022, before the peak of spring precipitation. Shelters were constructed using a metal frame extending to the outside of the perimeter of the plot that supported a 6 mm thick UV resistant clear plastic polyethylene film (VEVOR, Rancho Cucamonga, California, USA; (Fig. S1C). The metal frame was made of 0.75- and 1-inch diameter electrical metallic tubing conduit screwed to each other. It was elevated 1 m from the ground to allow ventilation, but it sat below the canopy of the trees to allow manipulation in the girdled plots (Fig. S1). The plastic was laid out such that it was possible to remove it and re-secure it for sampling. At this time, we also refreshed the trenches surrounding the plots and lined them with plastic to prevent re-connections.

This study is part of a larger project to examine, a) the carbon costs of osmoregulation (Sala et al. *in prep*), and b) potential conflicts between carbon allocation to storage-dependent water relations and to EcM fungi mediated via common ectomycorrhizal networks (this study). Due to logistical reasons, the EcM study was postponed until 2023 and the plastic from the rain shelters was therefore removed on September 24^th^ 2022, and reinstalled on June 2^nd^ 2023, to match the time in which we imposed girdling. Tyvek® was not reinstalled on the drought plots in the fall of 2022 to prevent excessive water stress in 2023. Trenches around the plots were revisited in the spring of 2023 to ensure they remained intact.

Girdling was implemented on June 2, 2023, by removing a 3-5 cm width band of bark and phloem approximately 50 cm above the ground around the trunk of the manipulated trees of girdle plots (5 ambient and 5 drought) (Fig. S1D). The girdled area was then covered with parafilm and reflective insulation to minimize exposure.

### Sampling and analyses

#### Osmotic, pressure, and water potential and NSC

To determine if osmotically active solutes or turgor were impacted by girdling neighboring trees and to account for variation among trees, we sampled R and T trees for predawn water potential (Ψ_pd_), osmotic potential, and NSC content of needles one week prior to girdling. We sampled needles from the 2021 needle cohort, which were produced before plots were set up, and therefore not impacted by the drought treatment for the separate experiment in 2022 (see above). Mid-canopy, south-facing terminal shoots were sampled at predawn (2 – 6 am) and transported to nearby measurement equipment. Within 5 minutes of collection, water potential was measured on a single fascicular bundle of 2-3 needles with a pressure chamber (PMS 1000, PMS Instrument Company, Albany, Oregon, USA)^69^. Simultaneously, approximately 5-10 bundles were wrapped tightly in labelled aluminum foil packets and frozen in liquid nitrogen to halt metabolic activity and minimize water loss. Frozen samples were transported on dry ice to the laboratory and stored at -20 °C until further osmotic potential and NSC measurements (see below). Additionally, punches of tree trunks (ca. 3 cm diameter) were taken from above and below the girdle in girdled trees and at approximately 50cm above the ground in response trees to assess the effects of the girdling treatment. Phloem tissue was separated from bark and wood and frozen in liquid nitrogen for further NSC analysis. These sampling procedures were repeated at 1, 2, 3, 4, 6, and 10 weeks following the initiation of drought and girdling treatments (Fig. 1).

Osmotic potential of needles was measured in the laboratory using a PSYPRO datalogger equipped with C-52 osmometer chambers (WESCOR, Inc, Logan, Utah, USA). One to three fascicles of frozen needles previously lysed in liquid nitrogen was used for sap extraction via centrifugation^70,71^. Needle tissue was packed with a ¼” diameter ball bearing in a 0.5 ml microcentrifuge tube and placed inside a larger 2 ml tube. A hole in the bottom of the smaller tube enabled collection of pressed sap into the larger tube when centrifuged at 10,000 RPM for 1 minute (Eppendorf 5415D, Eppendorf, Enfield Connecticut, USA).

Immediately following centrifugation, 10 μl of expressed sap was pipetted on to a filter paper disk and immediately sealed in the osmometer for measurement. Pressure potential was calculated as the difference between osmotic potential and water potential. All osmotic potential measurements were conducted within 4 days of field sampling to minimize the effect of sublimation in the freezer^72^.

To determine if soluble sugars and starch were also impacted by the girdling treatment of a neighbor tree, additional frozen needle and bole tissue from R and T trees were oven-dried at 70° C, coarsely ground with either a mortar and pestle (needles and phloem) or Wiley mill (xylem), and then finely ground to a homogeneous powder using a ball mill (TissueLyser II, QIAGEN, Germantown, Maryland, USA). Non-structural carbohydrate content was analyzed on approximately 12-14 mg of dry needle powder (or 10-12 mg of dry bole phloem powder) via enzymatic digestion to glucose and colorimetric quantification adapted from Sapes et al. (2021)^73^. Samples were incubated at 100 °C for one hour in 1.6 ml ultra-pure water to extract soluble sugars (glucose, fructose, and sucrose). Fructose and sucrose were converted into glucose using phosphoglucose isomerase (P5381, Sigma-Aldrich, St. Louis, MO, USA) and invertase (I4504, Sigma-Aldrich) and quantified with an ELx800 spectrometer (BioTek Instruments Inc, Winooski, VT, USA) at 340 nm after treated with hexokinase-glucose 6-phosphate dehydrogenase (G3293, Sigma-Aldrich). Total NSC concentration was obtained by digestion of polysaccharides (starch and sucrose) from a separate aliquot treated with amyloglucosidase (10115, Sigma-Aldrich) at 50 °C for 16 hours, followed by conversion to glucose and quantification as described above. Starch content was calculated as the difference between total NSC and soluble sugars. NSC components (soluble sugars, starch, and total NSC) are reported as a percentage of dry mass.

Volumetric soil water content was measured monthly at three random locations per plot with a 12 cm Hydrosense probe (Campbell Scientific, Logan Utah). Because the drought treatment started in the winter of 2021-2022, soil moisture was lower in drought plots (control and girdled) than in ambient plots before girdling and throughout the length of the study (Fig. S3).

#### Fungal biomass

To assess if the trenching depth was sufficient to sever most of the potential EcM connections, and to quantify any legacy of drought on fungal biomass prior to the girdling treatment, we sampled soil in 10 cm increments (2 cm Ø) to a depth of 40 cm in ambient and drought plots on May 20^th^, 2023. Soil was sieved (2 mm), placed in a 1.5 mL microcentrifuge tube, freeze-dried (Labconco Freezone benchtop freeze-dry system, Labconco, Kansas City, MO, USA) and stored at -20 °C prior to analyses. To quantify fungal biomass, we used phospholipid fatty acid (PLFA) analysis. This method is described in detail elsewhere^74^, but briefly, ∼2 g of freeze-dried soil was extracted with a chloroform-methanol-citrate buffer (1:2:0.8 v/v/v). We loaded extracted lipids onto silica gel columns (Bond Elut LRC, SI 100 mg; Varian, Palo Alto, California, USA), and the methanol fractions were subjected to mild alkaline methanolysis to form fatty acid methyl esters (FAMEs) prior to GC analyses. As there is no specific PLFA for EcM fungi, we used the PLFA 18:2ω6,9 to quantify all fungi^75^.

To evaluate responses in fungal biomass and EcM fungal communities (see below) to girdling and drought at the end of the experiment, three soil cores (2 cm Ø, 10 cm deep) were collected on September 30^th^, 2023, beneath the dripline of each tree and pooled for a total of 60 samples (20 plots x 3 trees per plot) and sieved and freeze-dried as before. Fatty acid analyses were conducted on pooled soil samples collected around Rc and C trees and from Rg and G trees (20 samples). We chose to pool soils within plots for fungal biomass analysis because we argue that the girdling effect would be present at the plot level, not necessarily isolated around individual trees as roots were distributed throughout the plot. Soil collected around the trenched trees were not included in biomass analyses as we expected trenching to isolate trenched trees from fungal dynamics associated with the girdled tree.

#### EcM fungal community analyses

The existence of common mycorrhizal networks among trees are exceedingly difficult to verify even with tracers. To address the potential for EcM networks to form, and to assess responses to various treatments, we used molecular analyses and compared EcM fungal community composition and richness in soil samples collected underneath individual trees at the end of the experiment in September 2023. Specifically, we compared EcM fungal communities collected beneath Rc and C trees in ambient and drought plots with the assumption that a high similarity in EcM fungal communities between samples would indicate potential for EcM networks to form. Likewise, we compared EcM fungal communities collected underneath Tc and Rc in ambient and drought plots to assess potential disturbance-related changes in fungal communities associated with trenching that could confound findings. Briefly, we used the Quick-DNA Fecal/Soil Microbe Miniprep kit (Zymo Research, Irvine, CA, USA) for DNA extraction from 250 mg of soil. Samples were prepared for Illumina sequencing through a two-step PCR amplification. We amplified the ITS2 region for general fungi using a mix of forward fungal primers flITS7 and flITS7o and the reverse primer ITS4^76^. Samples were assigned unique barcodes and pooled based on band strength using gel electrophoresis. Sequencing was performed at the University of Montana Genomics Core (umt.edu/genomics-lab/; Missoula, MT, USA). Amplicon libraries were sequenced using 2 x 250 paired end reads on the Illumina MiSeq platform (Illumnia Inc, San Diego, CA, USA). All sequences have been submitted to SRA on NCBI under accession number PRJNA1451872.

Raw sequence data was processed using “Quantitative Insights into Molecular Microbial Ecology 2” (Caporaso et al. 2010; QIIME2 version 2018.4). Forward and reverse reads were trimmed when median quality score fell below 30, which was 220bp for the forward reads and 180bp for the reverse reads. The q2-dada2 plugin was used to produce absolute sequence variants (ASVs) representing sequence clusters with 100% similarity. The database UNITE^77^ was used to assign taxonomy to representative fungal sequences. We then used the database FungalTraits^78,79^ to identify all taxa that matched with “ectomycorrhizal” under primary lifestyle, based on genus level assignments. For community analyses, we extracted all ASVs identified as “ectomycorrhizal” in soil and seedling root samples (see below) and rarified to 800 and 689 per sample, respectively, prior to statistical analysis using the q2-diversity plugin. At this sampling depth, rarefaction curves plateaued, which means we characterized the EcM fungal taxa that were in each sample. Three soil samples with fewer than 800 sequences were removed to allow for a greater sequencing depth. Additionally, the full fungal community was rarefied to 9,403 sequences, and the proportion of EcM fungal sequences per sample was compared with the assumption that if they did not differ, PLFA 18:2ω6,9 is a good proxy for EcM fungal biomass.

### Seedling experiment

To further assess the likelihood that responses in our experiment were due to EcM networks, we grew ponderosa pine seedlings in perforated cores within plots where connections to the EcM network can be manipulated by rotation^35^. We planted 5-month-old seedlings in May, 2023 (Fig. 1) into PVC cores (10 cm Ø, 25 cm deep) where 1 cm Ø holes had been drilled (∼30% of surface) and covered by a 31 μM mesh (Component Supply Company, Sparta, TN, USA) that allowed EcM fungal hyphae, but not roots, to enter. Seedlings had been reared in the lab from locally collected seeds grown in 100 ml conetainers (Stuewe and Sons, Tangent, OR, USA) filled with vermiculite and watered and fertilized with 20 ml of 0.5 g L^-1^ 20-2-20 (NPK, Peters Professional fertilizer, JR Peters Inc., Allentown, PA, USA) weekly to prevent nutrient limitations. Seedling roots were collected prior to transplanting and despite efforts to keep EcM fungi out of the lab, some root tips were colonized during rearing by a sequence variant within the genus *Pustularia*—a short-distance exploration type that does not form rhizomorphs (Põlme et al. 2020)—as determined from sequencing of the colonized root tips. This contamination did not preclude the use of seedlings as there were plenty of uncolonized root tips for other EcM fungi to colonize once outplanted, and molecular techniques allowed us to track the persistence of this contaminant in the field, which never exceeded 10%. The PVC cores in the field contained a 1:1 v:v mixture of sand:soil mix (NO_3_ ^-^: 3.6 mg kg soil^-1^, P :53 mg kg soil^-1^, K: 174 mg kg soil^-1^, pH 6.4). Soil was collected from the site and autoclaved to ensure that, upon transplanting to the field, EcM fungi other than the lab contaminant that colonized root tips came from outside the core and thus ensured that seedlings were connected to the network. Each core also included 0.5 tsp/L of hydrogels (The Scotts Company LLC., Marysville, OH, USA) for better water-holding capacity. Two cores per plot were planted on either side of the tree that would be girdled or not in the ambient treatment (rain shelters prevented us from planting cores in the droughted plots) and watered as needed for four months to allow colonization by EcM fungi (Fig. S2). By mid-September, rhizomorphs were visible based on observations from additional cores added for monitoring. On October 5^th^, 2023, we rotated one randomly chosen core per pair to rupture connections to the EcM network. At this time, we also girdled all additional trees, including control plots, to severely reduce C transfer from the trees to the EcM network (Fig. 1). We did not girdle trenched trees as they were assumed to be disconnected from the network. To assess if EcM fungi in the static cores extended beyond the core and EcM fungi in the rotated cores did not, we added 2 mg N as ^15^NH_415_NO_3_ (99% ^15^N) dissolved in 2.5 mL water, which we injected into 3 locations at 5-10 cm depth approximately 10 cm away from the core.

Eight days following girdling and ^15^N additions, seedlings were destructively harvested and divided into needles, stem, coarse roots (>2 mm Ø), and fine roots (including EcM root tips) for NSC analyses. To characterize the EcM fungal community pre- and post-transplanting, we randomly collected 3 root clusters with visible colonization from the root system. Those clusters were freeze-dried, ground, and then processed as outlined above. δ_15_N analyses were conducted by UC Davis Stable Isotope Facility (Davis, CA, USA) on ∼50 colonized root tips collected randomly per seedling using a dissecting scope and dried at 60° C. Colonized root tips from the unlabeled surplus cores were used for natural abundance of δ_15_N.

### Statistical Analyses

All analyses were conducted using R software v.4.5.0^80^. To assess the efficacy of the girdling treatment in reducing carbohydrate delivery to belowground, we assessed changes over time of soluble sugars, starch, and total NSC in the bole phloem sampled above, at, and below the girdle point in both C and G trees. We built linear mix effects models using package *lme4* for ambient and drought plots separately. Data for bole phloem starch, and total NSC were log-transformed to meet model assumptions. The fixed effect was the interaction between date and the variable defining the position relative to the gridle point (above, at, and below). Tree ID was used as a random effect. We considered effects to be significant if the 95% confidence interval of the estimates did not overlap with 0.

To evaluate the impacts of girdling on neighbor trees while accounting for environmental variability across space and time, we used a *difference in differences* (DiD) analytical approach^81,82^ using package *fixest*. In brief, DiD estimates the difference between a treated and untreated group for a given variable after treatment application, and relativizes that difference based on the existing difference between the two groups prior to applying the treatment. Since we estimate differences at each point in time and we use fixed and random effects, this specific kind of DiD is known as a *fixed effects event study*. Specifically, the response to girdling putatively mediated via the EcM network was assessed using the difference between responses in the R and T trees (R_G_ - T_G_) in the girdled plots, because EcM connections were severed in the T_G_ tree. This difference, however, also includes a trenching effect due to the disturbance created by trenching, which is measured in the control plots via the R_C_-T_C_ difference. Therefore, the response variable of our main DID model, (R_G_-T_G_) - (R_C_-T_C_), measures responses solely due to the putative EcM network while accounting for any potential side-effects caused by the trenching disturbance. The most critical assumption in this analysis is that any difference in pre-treatment means remains constant over time -known as parallel trends- so that they can be systematically subtracted to isolate the effects of treatment at each time step. In our case, there were no significant pre-treatment differences between control and treated groups, thus meeting the assumption. Because trade-offs are context dependent and depend on carbon supply and demand, we built separate sets of DIDs for ambient and drought plots. This allowed us to evaluate the strength of any potential trade-off between carbohydrate allocation to EcM and to storage-dependent osmoregulation under different supply-demand conditions. Dependent variables were leaf soluble sugars, leaf starch, leaf total NSC, predawn water potential, predawn osmotic potential, and predawn pressure potential. Treatment was defined as a dummy variable that considered if a tree was within a plot that received the girdling treatment or not.

The model is outlined in Equation 4. The sub-indexes t and d indicate tree and day, respectively. Dep_var represents each of the dependent variables; Manipulation_dummy*WeeksGirdling is an independent variable entered as a dummy that indicates whether values correspond to before (0) or after (1) girdling treatment. For DiD analyses of carbohydrate fractions (starch and soluble sugars) we also used Leaf water potential as an independent variable to account for potential confounding effects of water stress on carbohydrate fractions (Sapes et al. 2019). Tree_id and Days_girdling represent the random effects of tree id and days since girdling was applied.

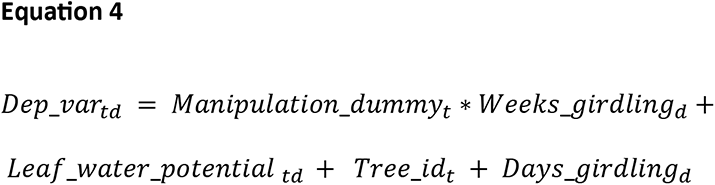

Finally, girdling times varied in the seedling study because the manipulated tree in girdle plots was girdled in June, and the remaining trees in all ambient plots (two trees in the control plots and response tree in the girdled plots) in October. To determine if this affected seedlings, we compared the total NSC content between control and girdled plots. Because we observed no differences (F _(1, 71)_ = 0.101, *p* = 0.764), we did not differentiate between control and girdled plots in downstream analyses.

### Fungal biomass and EcM fungal composition

Potential differences in fungal biomass over depth were assessed using a linear mixed-effects model with depth as a fixed effect and plot as a random effect. Linear mixed-effects models were also used to determine main and interactive effects between drought and girdling treatments on EcM fungal relative abundance and richness (obtained from sequence data) in soil samples with plot included as a random effect. Similar models were used to test whether δ¹⁵N and NSC differed between seedlings grown in static or rotated cores, with core treatment (static vs. rotated) and tissue type (fine root, coarse root, needle, and stem) as fixed effects and plot as a random effect. Violations of model assumptions were assessed using the DHARMa package. Data that did not meet model assumptions were log-transformed using the log1p function (natural log of *x* + 1) to improve normality and homoscedasticity.

To assess the potential for EcM network formation and to evaluate the effects of trenching, we analyzed pairwise differences in EcM fungal community composition using two models, soils collected adjacent to C and Rc trees, and soils collected adjacent to Rc and T trees, respectively. Specifically, we used perMANOVA on Bray-Curtis distances of Hellinger transformed rarefied sequence abundances using the *vegan* package and *adonis2* function^83^. We restricted both analyses to control plots as girdling could have affected outcomes. Tree ID, plot, and drought treatment were included as additive terms in the models to observe variation across plots and drought effects. In two separate models, perMANOVA was similarly used to 1) determine main and interactive effects of drought and girdling, where tree ID was nested within plot and to 2) assess if seedlings in static and rotated cores were colonized by different EcM fungal communities, where plot was included as an additive term.

Potential differences in EcM fungal communities were visualized using non-metric multidimensional scaling using the *metaMDS* function and plotted the results using the *ggplot2* package^84^. To identify individual EcM fungal taxa differentially abundant among treatments we used ANCOM^85^, which controls for false discovery rate and potential biases associated with compositional data^86^. For this analysis, we used non-rarefied sequence data that included all fungi.

## Acknowledgements

This project was funded by the Department of Energy (project DE-FOA-0002392) and by MPG Ranch. This research was supported by University of Montana Genomics Core and Montana INBRE Data Science Core, which are funded by the National Institute of General Medical Sciences (P20GM103474), the Office of the Vice President for Research and Creative Scholarship at the University of Montana, and the M. J. Murdock Charitable Trust. The content is solely the responsibility of the authors and does not necessarily represent the official views of the UMGC or the National Institutes of Health. We thank The Nature Conservancy and Paws Up Ranch for allowing us to perform this study on their property. Jake Kleimann, Jazmine Raymond, Ella Keefer, Sarah Tremper, Nora Christie, and McKenzie Murphy helped with field data collection. We also thank J. Karst and anonymous reviewers for their comments in previous versions of this manuscript.

## Author contributions

A.S., Y.L., R.K., and G.S. conceived the study, A.S., Y.L., R.K., and G.S. obtained funding, A.G., M.E.D., A.S., and Y.L. gathered and prepared the data, A.G., G.S., M.E.D., Y.L., and L.B. designed and developed the analyses, G.S., M.E.D., and Y.L. prepared the results and figures, G.S., A.S., M.E.D, and Y.L. wrote the manuscript with contributions of A.G., L.B., and R.K..

## Competing interests

The authors declare no competing interests.

Supplementary Information is available for this paper.

Correspondence and requests for materials should be addressed to Gerard Sapes.

