## Supplementary Information for "Ectomycorrhizal networks propagate carbon deficit and drought stress among trees"

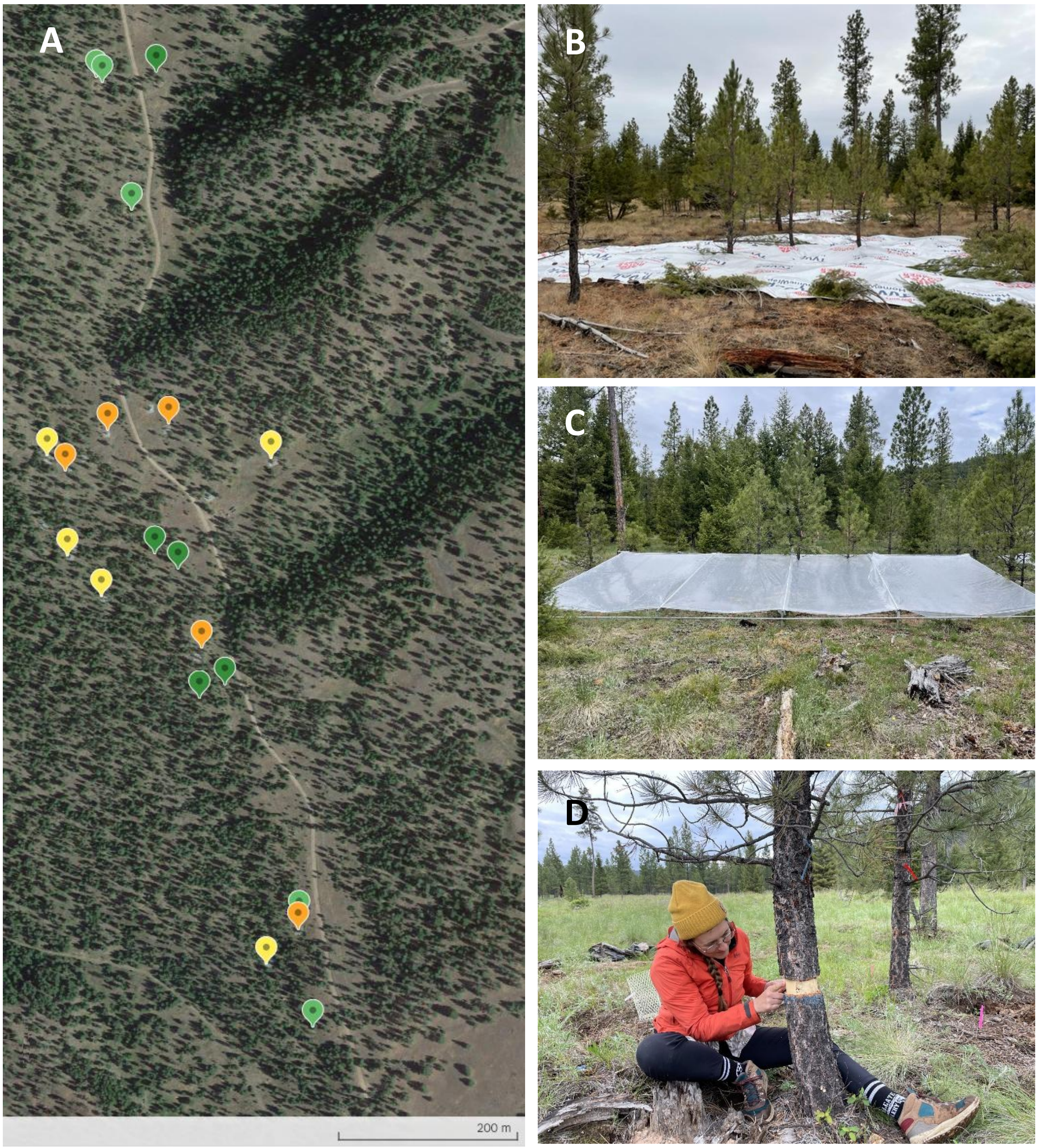


**Figure S1:** A) Aerial view of the study site with the drought-control (yellow), drought-girdled (orange), ambient-control (light green), and ambient-girdled (dark green) plots. B) Example of a Tyvek® installation to exclude snow precipitation in winter 2021. C) Example of a rain shelter installation to exclude precipitation. D) Example of girdling of one of the manipulated trees.

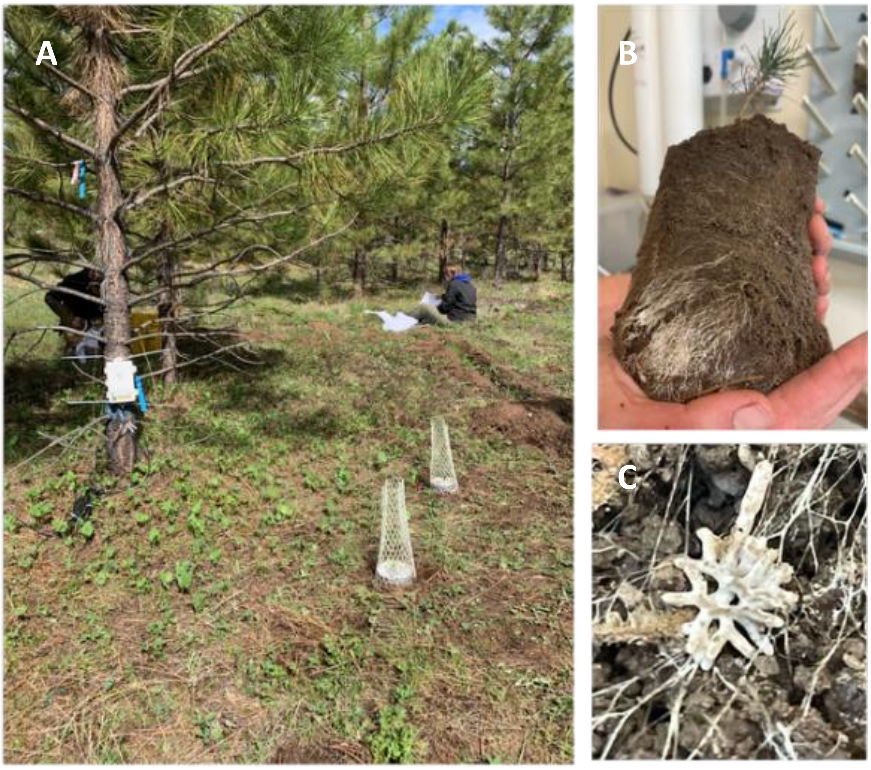


**Figure S2:** Seedling experiment where one out of the two cores (a) was rotated after 5 months following visual colonization by surrounding EcMF fungi (b), including rhizomorphs (c).

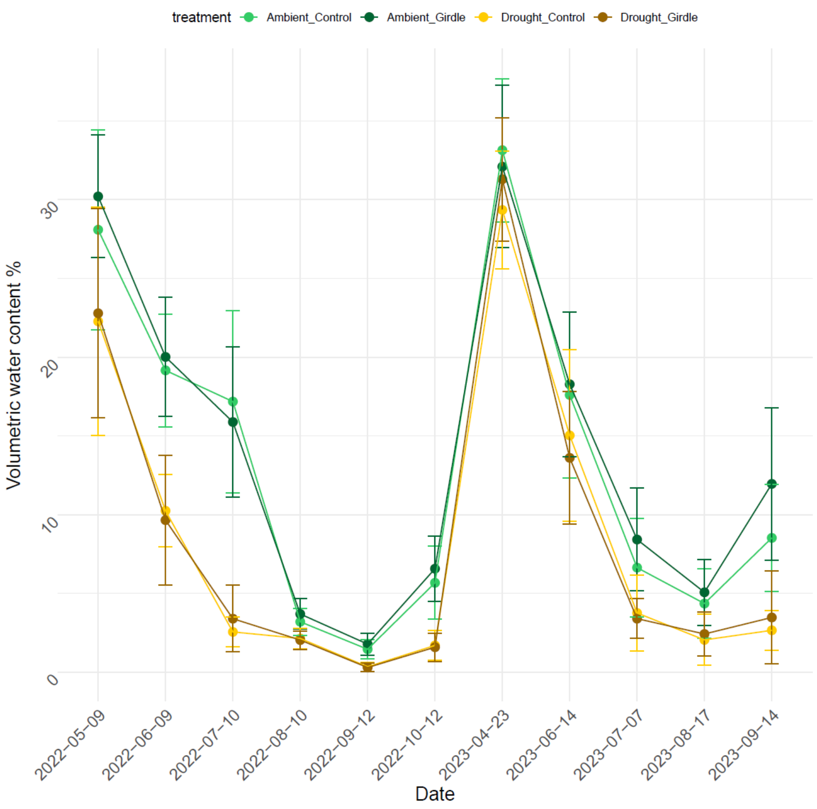


```
Call:
lm(formula = SoilVolumetricWaterContent ~ Treatment * Date, data = vwc)

Residuals:
     Min       1Q   Median       3Q      Max 
-17.3500  -1.3844  -0.1272   1.2731  20.2544 

Coefficients:
                                    Estimate Std. Error t value Pr(>|t|)    
(Intercept)                          29.1500     0.3722  78.310   < 2e-16 ***
TreatmentDrought                     -6.6156     0.5264 -12.567   < 2e-16 ***
Date2022-06-09                       -9.5567     0.5264 -18.154   < 2e-16 ***
Date2022-07-10                      -12.6144     0.5264 -23.962   < 2e-16 ***
Date2022-08-10                      -25.6967     0.5264 -48.813   < 2e-16 ***
Date2022-09-12                      -27.5244     0.5264 -52.285   < 2e-16 ***
Date2022-10-12                      -23.0222     0.5264 -43.733   < 2e-16 ***
Date2023-04-23                        3.4711     0.5264   6.594  5.50e-11 ***
Date2023-06-14                      -11.2044     0.5264 -21.284   < 2e-16 ***
Date2023-07-07                      -21.6156     0.5264 -41.061   < 2e-16 ***
Date2023-08-17                      -24.4289     0.5264 -46.405   < 2e-16 ***
Date2023-09-14                      -18.9044     0.5264 -35.911   < 2e-16 ***
TreatmentDrought:Date2022-06-09     -3.0211     0.7445  -4.058  5.14e-05 ***
TreatmentDrought:Date2022-07-10     -6.9378     0.7445  -9.319   < 2e-16 ***
TreatmentDrought:Date2022-08-10      5.2589     0.7445   7.064  2.24e-12 ***
TreatmentDrought:Date2022-09-12      5.3167     0.7445   7.141  1.30e-12 ***
TreatmentDrought:Date2022-10-12      2.1356     0.7445   2.869  0.004168 ** 
TreatmentDrought:Date2023-04-23      4.3178     0.7445   5.800  7.73e-09 ***
TreatmentDrought:Date2023-06-14      2.9941     0.7445   4.022  6.00e-05 ***
TreatmentDrought:Date2023-07-07      2.6656     0.7445   3.580  0.000351 ***
TreatmentDrought:Date2023-08-17      4.1378     0.7445   5.558  3.10e-08 ***
TreatmentDrought:Date2023-09-14     -0.5578     0.7445  -0.749  0.453816    
---
Signif. codes:  0 ‘***’ 0.001 ‘**’ 0.01 ‘*’ 0.05 ‘.’ 0.1 ‘ ’ 1

Residual standard error: 3.531 on 1958 degrees of freedom
Multiple R-squared:  0.8915, Adjusted R-squared:  0.8903 
F-statistic: 765.8 on 21 and 1958 DF,  p-value: < 2.2e-16
```

**Figure S3:** Dynamics of soil volumetric water content in ambient control (light green), ambient girdle (dark green), drought control (yellow), and drought girdle (brown) throughout the experiment.

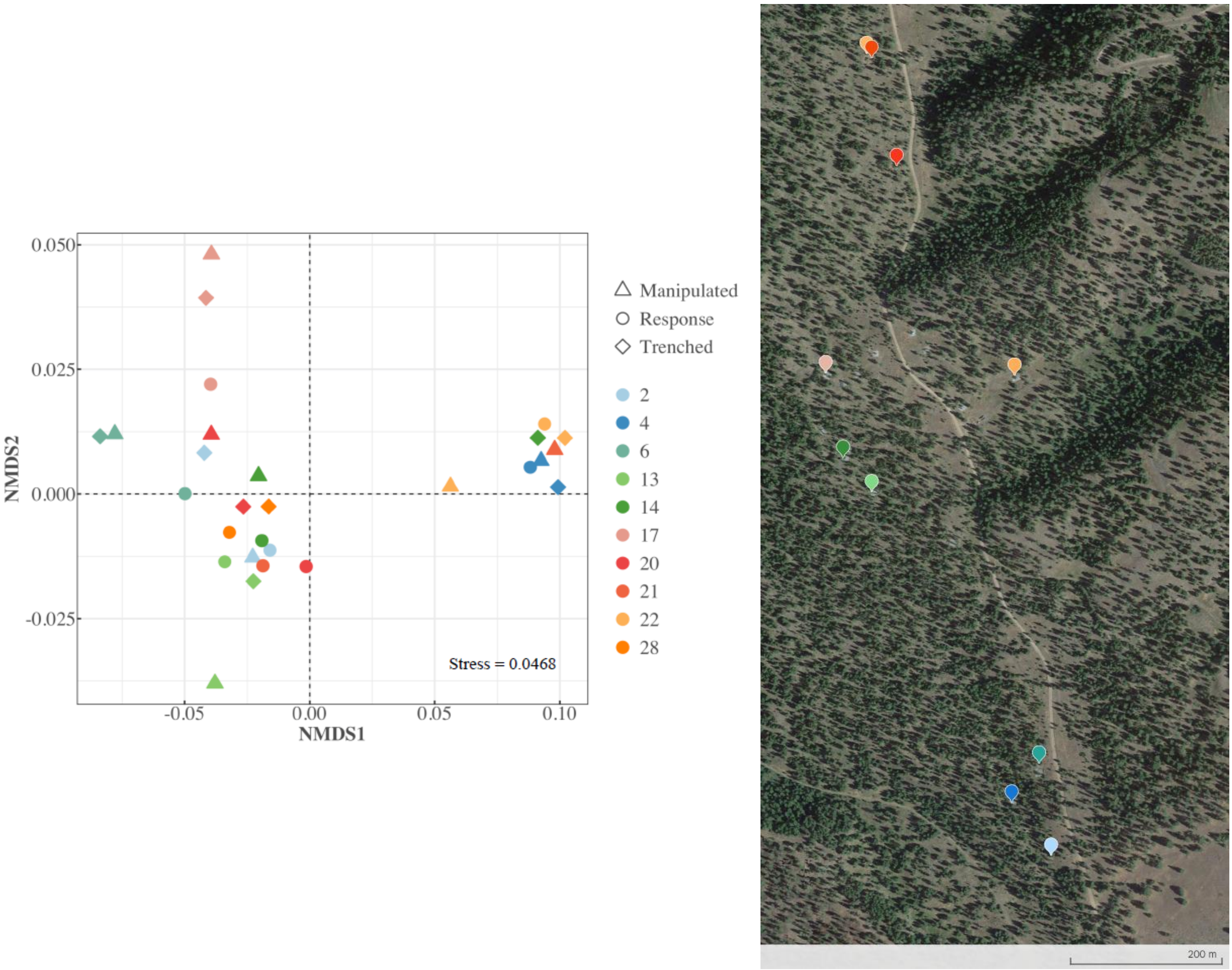


**Figure S4:** NMDS of EcM fungal community composition across plots and tree assignment in the control plots and map with the location of the plots across space. Plots are color coded by proximity to highlight the substantially high variability in fungal communities among neighboring plots relative to within plots.

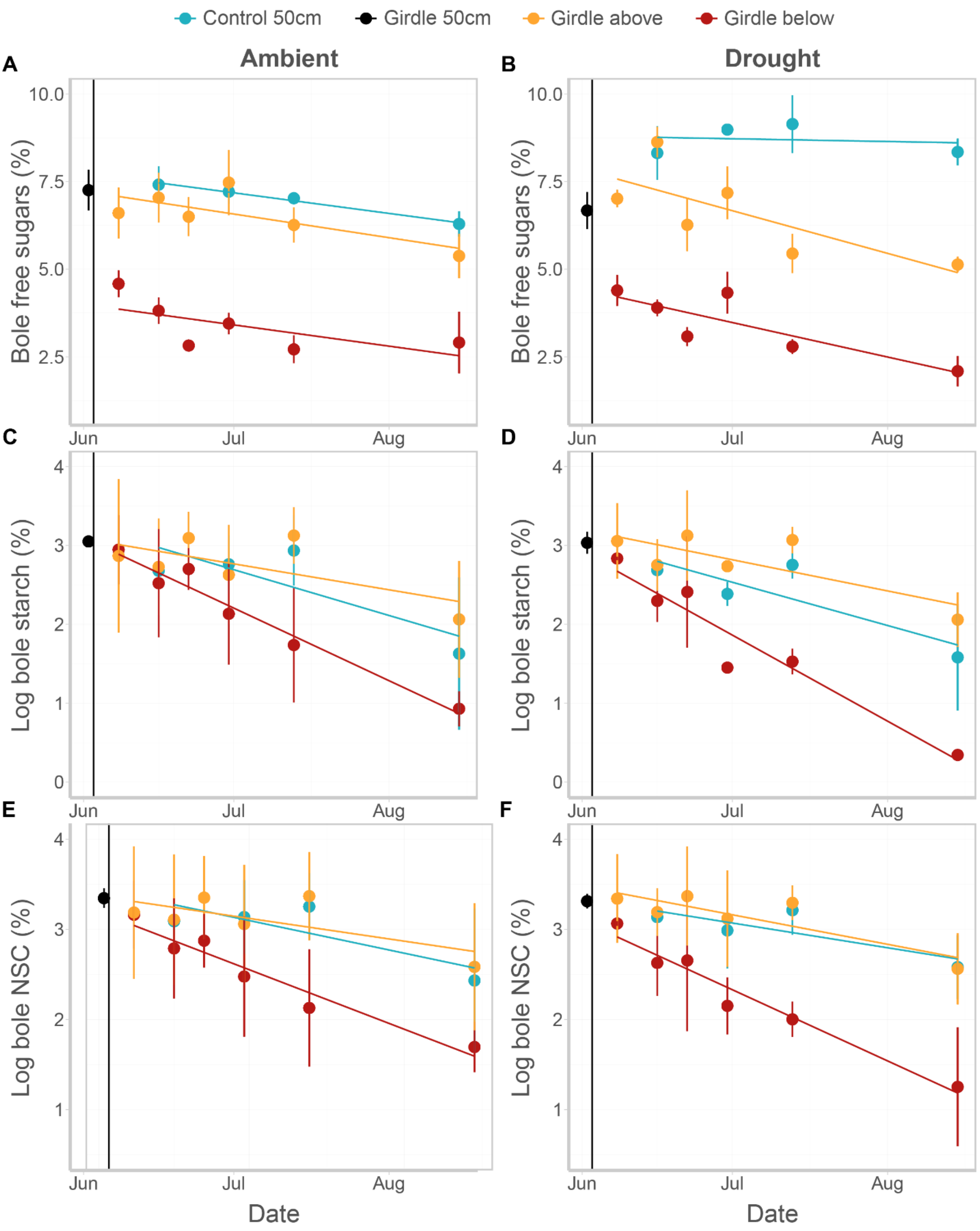


**Figure S5:** Girdling caused NSC depletion below the wound (red) relative to above the wound (orange) by impairing transfer of NSCs towards belowground sinks. Concentrations above the would were similar to concentrations found in control (i.e., non-girdled) trees (blue).

**Table S1:** Linear mix effects models explaining carbohydrate dynamics in manipulated trees in response to girdling. Blue highlights factors with estimate confidence intervals not overlapping with 0.

| | | | | | | |
|---|---|---|---|---|---|---|
| **Ambient** | | | | | | |
| **A** | | | | | | |
| **Soluble sugars** | | | | | | |
| **Model and factors** | **Model type** | **Estimate** | **2.5% C.I.** | **97.5% C.I.** | **T-value** | |
| *BolePhloemSolubleSugars ~ DaysSinceGirdling * ManipulationxPosition + 1 \| TreeTag* | LMER | | | | | |
| *sd_Intercept\|TreeTag* | | | 0.000 | 0.688 | | |
| *sigma* | | 1.158 | 0.961 | 1.331 | | |
| *Intercept* | | 7.728 | 6.689 | 8.767 | 14.133 | |
| *DaysSinceGirdling* | | -0.019 | -0.041 | 0.003 | -1.626 | |
| *ManipulationxPositionGirdle.above* | | -0.523 | -1.794 | 0.748 | -0.780 | |
| *ManipulationxPositionGirdle.below* | | -3.750 | -5.021 | -2.479 | -5.590 | * |
| *DaysSinceGirdling x ManipulationxPositionGirdle.above* | | -0.003 | -0.032 | 0.026 | -0.188 | |
| *DaysSinceGirdling x ManipulationxPositionGirdle.below* | | -0.001 | -0.029 | 0.028 | -0.040 | |
| **B** | | | | | | |
| **Starch** | | | | | | |
| **Model and factors** | **Model type** | **Estimate** | **2.5% C.I.** | **97.5% C.I.** | **T-value** | |
| *logBolePhloemStarch ~ DaysSinceGirdling * ManipulationxPosition + 1 \| TreeTag* | LMER | | | | | |
| *sd_Intercept\|TreeTag* | | | 0.000 | 0.285 | | |
| *sigma* | | 0.450 | 0.373 | 0.516 | | |
| *Intercept* | | 3.233 | 2.824 | 3.641 | 15.030 | |
| *DaysSinceGirdling* | | -0.019 | -0.028 | -0.010 | -4.169 | * |
| *ManipulationxPositionGirdle.above* | | -0.201 | -0.703 | 0.300 | -0.760 | |
| *ManipulationxPositionGirdle.below* | | -0.133 | -0.635 | 0.368 | -0.504 | |
| *DaysSinceGirdling x ManipulationxPositionGirdle.above* | | 0.009 | -0.002 | 0.020 | 1.515 | |
| *DaysSinceGirdling x ManipulationxPositionGirdle.below* | | -0.016 | -0.028 | -0.005 | -2.798 | * |
| **C** | | | | | | |
| **Total NSC** | | | | | | |
| **Model and factors** | **Model type** | **Estimate** | **2.5% C.I.** | **97.5% C.I.** | **T-value** | |
| *logBolePhloemTotalNSC ~ DaysSinceGirdling * ManipulationxPosition + 1 \| TreeTag* | LMER | | | | | |
| *sd_Intercept\|TreeTag* | | | 0.000 | 0.253 | | |
| *sigma* | | 0.390 | 0.324 | 0.448 | | |
| *Intercept* | | 3.433 | 3.078 | 3.789 | 18.322 | |
| *DaysSinceGirdling* | | -0.012 | -0.019 | -0.004 | -2.981 | * |
| *ManipulationxPositionGirdle.above* | | -0.093 | -0.531 | 0.345 | -0.402 | |
| *ManipulationxPositionGirdle.below* | | -0.182 | -0.619 | 0.256 | -0.786 | |
| *DaysSinceGirdling x ManipulationxPositionGirdle.above* | | 0.004 | -0.006 | 0.013 | 0.741 | |
| *DaysSinceGirdling x ManipulationxPositionGirdle.below* | | -0.016 | -0.025 | -0.006 | -3.093 | * |

**Table S1 (continued)**

| Drought | | | | | | |
|---|---|---|---|---|---|---|
| **D** | | | | | | |
| **Soluble sugars** | | | | | | |
| **Model and factors** | **Model type** | **Estimate** | **2.5% C.I.** | **97.5% C.I.** | **T-value** | |
| *BolePhloemSolubleSugars ~ DaysSinceGirdling * ManipulationxPosition + 1 \| TreeTag* | LMER | | | | | |
| *sd_Intercept\|TreeTag* | | | 0.000 | 1.306 | | |
| *sigma* | | 1.230 | 1.021 | 1.443 | | |
| *Intercept* | | 8.795 | 7.574 | 10.016 | 13.463 | |
| *DaysSinceGirdling* | | -0.003 | -0.027 | 0.021 | -0.206 | |
| *ManipulationxPositionGirdle.above* | | -0.988 | -2.533 | 0.558 | -1.192 | |
| *ManipulationxPositionGirdle.below* | | -4.396 | -5.942 | -2.851 | -5.307 | * |
| *DaysSinceGirdling x ManipulationxPositionGirdle.above* | | -0.037 | -0.068 | -0.006 | -2.302 | * |
| *DaysSinceGirdling x ManipulationxPositionGirdle.below* | | -0.029 | -0.060 | 0.002 | -1.830 | |
| **E** | | | | | | |
| **Starch** | | | | | | |
| **Model and factors** | **Model type** | **Estimate** | **2.5% C.I.** | **97.5% C.I.** | **T-value** | |
| *logBolePhloemStarch ~ DaysSinceGirdling * ManipulationxPosition + 1 \| TreeTag* | LMER | | | | | |
| *sd_Intercept\|TreeTag* | | | 0.000 | 0.223 | | |
| *sigma* | | 0.383 | 0.318 | 0.440 | | |
| *Intercept* | | 3.031 | 2.688 | 3.375 | 16.771 | |
| *DaysSinceGirdling* | | -0.018 | -0.025 | -0.010 | -4.601 | * |
| *ManipulationxPositionGirdle.above* | | 0.166 | -0.255 | 0.586 | 0.748 | |
| *ManipulationxPositionGirdle.below* | | -0.220 | -0.640 | 0.200 | -0.993 | |
| *DaysSinceGirdling x ManipulationxPositionGirdle.above* | | 0.004 | -0.005 | 0.014 | 0.821 | |
| *DaysSinceGirdling x ManipulationxPositionGirdle.below* | | -0.017 | -0.027 | -0.008 | -3.521 | * |
| **F** | | | | | | |
| **Total NSC** | | | | | | |
| **Model and factors** | **Model type** | **Estimate** | **2.5% C.I.** | **97.5% C.I.** | **T-value** | |
| *logBolePhloemTotalNSC ~ DaysSinceGirdling * ManipulationxPosition + 1 \| TreeTag* | LMER | | | | | |
| *sd_Intercept\|TreeTag* | | | 0.000 | 0.159 | | |
| *sigma* | | 0.253 | 0.210 | 0.290 | | |
| *Intercept* | | 3.319 | 3.090 | 3.547 | 27.550 | |
| *DaysSinceGirdling* | | -0.009 | -0.014 | -0.004 | -3.470 | * |
| *ManipulationxPositionGirdle.above* | | 0.149 | -0.132 | 0.430 | 1.006 | |
| *ManipulationxPositionGirdle.below* | | -0.272 | -0.553 | 0.009 | -1.836 | |
| *DaysSinceGirdling x ManipulationxPositionGirdle.above* | | -0.002 | -0.008 | 0.004 | -0.629 | |
| *DaysSinceGirdling x ManipulationxPositionGirdle.below* | | -0.017 | -0.024 | -0.011 | -5.285 | * |

**Table S2:** Linear mix effects models comparing rates of carbohydrate decline in manipulated trees between ambient and drought plots. Blue highlights factors with estimate confidence intervals not overlapping with 0.

**Ambient & Drought**

**A**

**Soluble sugars**

| Model and factors | Model type | Estimate | 2.5% C.I. | 97.5% C.I. | T-value | |
|---|---|---|---|---|---|---|
| *BolePhloemSolubleSugars ~ DaysSinceGirdling * TreatmentxManipulationxPosition + 1 \| TreeTag* | LMER | | | | | |
| *sd_Intercept\|TreeTag* | | | 0 | 0.58781 | | |
| *sigma* | | 0.98952 | 0.8066 | 1.18416 | | |
| *Intercept* | | 3.97844 | 3.36311 | 4.59377 | 12.240017 | |
| *DaysSinceGirdling* | | -0.01956 | -0.0354 | -0.00371 | -2.4169789 | * |
| *TreatmentxManipulationxPositionDrought.Girdle.below* | | 0.42011 | -0.4501 | 1.29032 | 0.9139415 | |
| *DaysSinceGirdling x TreatmentxManipulationxPositionDrought.Girdle.below* | | -0.01218 | -0.03458 | 0.01022 | -1.0643514 | |

**B**

**Starch**

| Model and factors | Model type | Estimate | 2.5% C.I. | 97.5% C.I. | T-value | |
|---|---|---|---|---|---|---|
| *logBolePhloemStarch ~ DaysSinceGirdling * TreatmentxManipulationxPosition + 1 \| TreeTag* | LMER | | | | | |
| *sd_Intercept\|TreeTag* | | | 0.14811 | 0.5597 | | |
| *sigma* | | 0.45655 | 0.37215 | 0.55167 | | |
| *Intercept* | | 3.09919 | 2.70132 | 3.49706 | 14.77167 | |
| *DaysSinceGirdling* | | -0.03519 | -0.0425 | -0.02788 | -9.4260627 | * |
| *TreatmentxManipulationxPositionDrought.Girdle.below* | | -0.28818 | -0.85085 | 0.2745 | -0.9712419 | |
| *DaysSinceGirdling x TreatmentxManipulationxPositionDrought.Girdle.below* | | -3.00E-05 | -0.01036 | 0.01031 | -0.0050966 | |

**C**

**Total NSC**

| Model and factors | Model type | Estimate | 2.5% C.I. | 97.5% C.I. | T-value | |
|---|---|---|---|---|---|---|
| *logBolePhloemTotalNSC ~ DaysSinceGirdling * TreatmentxManipulationxPosition + 1 \| TreeTag* | LMER | | | | | |
| *sd_Intercept\|TreeTag* | | | 0.10775 | 0.46643 | | |
| *sigma* | | 0.40935 | 0.33368 | 0.49464 | | |
| *Intercept* | | 3.25168 | 2.91117 | 3.59219 | 18.089427 | |
| *DaysSinceGirdling* | | -0.02734 | -0.03389 | -0.02079 | -8.1688261 | * |
| *TreatmentxManipulationxPositionDrought.Girdle.below* | | -0.20524 | -0.6868 | 0.27631 | -0.807369 | |
| *DaysSinceGirdling x TreatmentxManipulationxPositionDrought.Girdle.below* | | 0.0012 | -0.00807 | 0.01047 | 0.2536149 | |

**Table S3:** Difference in differences models assessing changes in water relations variables when accounting for both response and trenched trees in ambient and drought plots. Bold highlights indicate significant effects. Significance levels: · = p < 0.1, * = p < 0.05.

**Ambient**

**A**

**Pressure potential**

| Factors | Event | Estimate | Std. error | P-value | Sig. | Adj. R2 | 2.5% C.I. | 97.5% C.I. | Days girdling |
|---|---|---|---|---|---|---|---|---|---|
| *M_dummy * WeeksGirdling::0* | 0 | 0.000 | 0.000 | | | | 0.000 | 0.000 | -3 |
| *M_dummy * WeeksGirdling::1* | 1 | -0.192 | 0.171 | 0.292 | | | -0.580 | 0.196 | 6 |
| *M_dummy * WeeksGirdling::2* | 2 | -0.028 | 0.096 | 0.777 | | | -0.245 | 0.189 | 14 |
| *M_dummy * WeeksGirdling::3* | 3 | -0.032 | 0.108 | 0.774 | | 0.324 | -0.276 | 0.212 | 20 |
| *M_dummy * WeeksGirdling::4* | 4 | -0.154 | 0.092 | 0.127 | | | -0.361 | 0.053 | 28 |
| *M_dummy * WeeksGirdling::5* | 5 | -0.162 | 0.165 | 0.352 | | | -0.535 | 0.211 | 41 |
| *M_dummy * WeeksGirdling::6* | 6 | -0.022 | 0.101 | 0.832 | | | -0.250 | 0.206 | 74 |

**B**

**Osmotic potential**

| Factors | Event | Estimate | Std. error | P-value | Sig. | Adj. R2 | 2.5% C.I. | 97.5% C.I. | Days girdling |
|---|---|---|---|---|---|---|---|---|---|
| *M_dummy * WeeksGirdling::0* | 0 | 0.000 | 0.000 | | | | 0.000 | 0.000 | -3 |
| *M_dummy * WeeksGirdling::1* | 1 | 0.123 | 0.153 | 0.441 | | | -0.223 | 0.469 | 6 |
| *M_dummy * WeeksGirdling::2* | 2 | -0.085 | 0.114 | 0.479 | | | -0.343 | 0.174 | 14 |
| *M_dummy * WeeksGirdling::3* | 3 | -0.018 | 0.133 | 0.895 | | 0.494 | -0.318 | 0.282 | 20 |
| *M_dummy * WeeksGirdling::4* | 4 | 0.004 | 0.103 | 0.971 | | | -0.229 | 0.237 | 28 |
| *M_dummy * WeeksGirdling::5* | 5 | 0.047 | 0.107 | 0.670 | | | -0.195 | 0.289 | 41 |
| *M_dummy * WeeksGirdling::6* | 6 | -0.048 | 0.123 | 0.704 | | | -0.325 | 0.229 | 74 |

**C**

**Water potential**

| Factors | Event | Estimate | Std. error | P-value | Sig. | Adj. R2 | 2.5% C.I. | 97.5% C.I. | Days girdling |
|---|---|---|---|---|---|---|---|---|---|
| *M_dummy * WeeksGirdling::0* | 0 | 0.000 | 0.000 | | | | 0.000 | 0.000 | -3 |
| *M_dummy * WeeksGirdling::1* | 1 | -0.070 | 0.058 | 0.260 | | | -0.202 | 0.062 | 6 |
| *M_dummy * WeeksGirdling::2* | 2 | -0.110 | 0.066 | 0.132 | | | -0.260 | 0.040 | 14 |
| *M_dummy * WeeksGirdling::3* | 3 | -0.050 | 0.073 | 0.508 | | 0.250 | -0.214 | 0.114 | 20 |
| *M_dummy * WeeksGirdling::4* | 4 | -0.130 | 0.087 | 0.169 | | | -0.327 | 0.067 | 28 |
| *M_dummy * WeeksGirdling::5* | 5 | -0.120 | 0.116 | 0.329 | | | -0.383 | 0.143 | 41 |
| *M_dummy * WeeksGirdling::6* | 6 | -0.070 | 0.090 | 0.458 | | | -0.274 | 0.134 | 74 |

### Table S3 (continued)

**Drought**

**D**

**Pressure potential**

| Factors | Event | Estimate | Std. error | P-value | Sig. | Adj. R2 | 2.5% C.I. | 97.5% C.I. | Days girdling |
|---|---|---|---|---|---|---|---|---|---|
| *M_dummy * WeeksGirdling::0* | 0 | 0.000 | 0.000 | | | | 0.000 | 0.000 | -3 |
| ***M_dummy * WeeksGirdling::1*** | **1** | **-0.250** | **0.105** | **0.041** | ***** | | **-0.488** | **-0.012** | **6** |
| *M_dummy * WeeksGirdling::2* | 2 | -0.011 | 0.096 | 0.911 | | | -0.228 | 0.206 | 14 |
| *M_dummy * WeeksGirdling::3* | 3 | -0.236 | 0.161 | 0.177 | | 0.256 | -0.600 | 0.128 | 20 |
| ***M_dummy * WeeksGirdling::4*** | **4** | **-0.174** | **0.078** | **0.052** | **.** | | **-0.350** | **0.002** | **28** |
| *M_dummy * WeeksGirdling::5* | 5 | -0.176 | 0.122 | 0.184 | | | -0.453 | 0.101 | 41 |
| *M_dummy * WeeksGirdling::6* | 6 | -0.135 | 0.115 | 0.270 | | | -0.395 | 0.125 | 74 |

**E**

**Osmotic potential**

| Factors | Event | Estimate | Std. error | P-value | Sig. | Adj. R2 | 2.5% C.I. | 97.5% C.I. | Days girdling |
|---|---|---|---|---|---|---|---|---|---|
| *M_dummy * WeeksGirdling::0* | 0 | 0.000 | 0.000 | | | | 0.000 | 0.000 | -3 |
| *M_dummy * WeeksGirdling::1* | 1 | 0.096 | 0.120 | 0.443 | | | -0.176 | 0.368 | 6 |
| *M_dummy * WeeksGirdling::2* | 2 | -0.005 | 0.124 | 0.967 | | | -0.287 | 0.276 | 14 |
| *M_dummy * WeeksGirdling::3* | 3 | 0.191 | 0.136 | 0.194 | | 0.317 | -0.117 | 0.500 | 20 |
| *M_dummy * WeeksGirdling::4* | 4 | 0.100 | 0.085 | 0.267 | | | -0.091 | 0.292 | 28 |
| *M_dummy * WeeksGirdling::5* | 5 | 0.012 | 0.131 | 0.932 | | | -0.284 | 0.307 | 41 |
| *M_dummy * WeeksGirdling::6* | 6 | 0.017 | 0.106 | 0.876 | | | -0.223 | 0.257 | 74 |

**F**

**Water potential**

| Factors | Event | Estimate | Std. error | P-value | Sig. | Adj. R2 | 2.5% C.I. | 97.5% C.I. | Days girdling |
|---|---|---|---|---|---|---|---|---|---|
| *M_dummy * WeeksGirdling::0* | 0 | 0.000 | 0.000 | | | | 0.000 | 0.000 | -3 |
| ***M_dummy * WeeksGirdling::1*** | **1** | **-0.160** | **0.078** | **0.071** | **.** | | **-0.337** | **0.017** | **6** |
| *M_dummy * WeeksGirdling::2* | 2 | -0.015 | 0.066 | 0.825 | | | -0.164 | 0.134 | 14 |
| *M_dummy * WeeksGirdling::3* | 3 | -0.060 | 0.090 | 0.520 | | 0.405 | -0.263 | 0.143 | 20 |
| *M_dummy * WeeksGirdling::4* | 4 | -0.080 | 0.066 | 0.257 | | | -0.230 | 0.070 | 28 |
| *M_dummy * WeeksGirdling::5* | 5 | -0.190 | 0.109 | 0.116 | | | -0.437 | 0.057 | 41 |
| *M_dummy * WeeksGirdling::6* | 6 | -0.095 | 0.180 | 0.611 | | | -0.503 | 0.313 | 74 |

**Table S4:** Difference in differences models assessing changes in non-structural carbohydrate variables when accounting for both response and trenched trees in ambient and drought plots. Bold highlights indicate significant effects. Significance levels: * = p < 0.05.

**Ambient**

**A**

**Soluble sugars**

| Factors | Event | Estimate | Std. error | P-value | Sig. | Adj. R2 | 2.5% C.I. | 97.5% C.I. | Days girdling |
|---|---|---|---|---|---|---|---|---|---|
| *M_dummy * WeeksGirdling::0* | 0 | 0.000 | 0.000 | | | | 0.000 | 0.000 | -3 |
| *M_dummy * WeeksGirdling::1* | 1 | 0.206 | 0.626 | 0.750 | | | -1.209 | 1.621 | 6 |
| *M_dummy * WeeksGirdling::2* | 2 | -0.436 | 0.358 | 0.254 | | | -1.245 | 0.373 | 14 |
| *M_dummy * WeeksGirdling::3* | 3 | 0.028 | 0.429 | 0.949 | | 0.735 | -0.943 | 0.999 | 20 |
| *M_dummy * WeeksGirdling::4* | 4 | -0.082 | 0.587 | 0.892 | | | -1.410 | 1.245 | 28 |
| *M_dummy * WeeksGirdling::5* | 5 | -0.398 | 0.313 | 0.237 | | | -1.107 | 0.312 | 41 |
| *M_dummy * WeeksGirdling::6* | 6 | 0.275 | 0.418 | 0.527 | | | -0.671 | 1.222 | 74 |

**B**

**Starch**

| Factors | Event | Estimate | Std. error | P-value | Sig. | Adj. R2 | 2.5% C.I. | 97.5% C.I. | Days girdling |
|---|---|---|---|---|---|---|---|---|---|
| *M_dummy * WeeksGirdling::0* | 0 | 0.000 | 0.000 | | | | 0.000 | 0.000 | -3 |
| *M_dummy * WeeksGirdling::1* | 1 | -1.351 | 2.343 | 0.578 | | | -6.652 | 3.950 | 6 |
| *M_dummy * WeeksGirdling::2* | 2 | -0.487 | 1.773 | 0.790 | | | -4.498 | 3.523 | 14 |
| *M_dummy * WeeksGirdling::3* | 3 | -1.910 | 2.124 | 0.392 | | 0.500 | -6.715 | 2.895 | 20 |
| *M_dummy * WeeksGirdling::4* | 4 | 0.192 | 1.787 | 0.917 | | | -3.851 | 4.234 | 28 |
| *M_dummy * WeeksGirdling::5* | 5 | -1.560 | 1.910 | 0.435 | | | -5.881 | 2.760 | 41 |
| *M_dummy * WeeksGirdling::6* | 6 | -3.011 | 2.192 | 0.203 | | | -7.970 | 1.947 | 74 |

**C**

**Total NSC**

| Factors | Event | Estimate | Std. error | P-value | Sig. | Adj. R2 | 2.5% C.I. | 97.5% C.I. | Days girdling |
|---|---|---|---|---|---|---|---|---|---|
| *M_dummy * WeeksGirdling::0* | 0 | 0.000 | 0.000 | | | | 0.000 | 0.000 | -3 |
| *M_dummy * WeeksGirdling::1* | 1 | -1.148 | 2.193 | 0.613 | | | -6.110 | 3.813 | 6 |
| *M_dummy * WeeksGirdling::2* | 2 | -0.917 | 1.742 | 0.611 | | | -4.858 | 3.024 | 14 |
| *M_dummy * WeeksGirdling::3* | 3 | -1.881 | 1.948 | 0.359 | | 0.598 | -6.287 | 2.525 | 20 |
| *M_dummy * WeeksGirdling::4* | 4 | 0.158 | 1.621 | 0.924 | | | -3.508 | 3.825 | 28 |
| *M_dummy * WeeksGirdling::5* | 5 | -1.970 | 1.994 | 0.349 | | | -6.482 | 2.542 | 41 |
| *M_dummy * WeeksGirdling::6* | 6 | -2.736 | 2.294 | 0.264 | | | -7.926 | 2.454 | 74 |

### Table S4 (continued)

**Drought**

**D**

**Soluble sugars**

| Factors | Event | Estimate | Std. error | P-value | Sig. | Adj. R2 | 2.5% C.I. | 97.5% C.I. | Days girdling |
|---|---|---|---|---|---|---|---|---|---|
| *M_dummy * WeeksGirdling::0* | 0 | 0.000 | 0.000 | | | | 0.000 | 0.000 | -3 |
| *M_dummy * WeeksGirdling::1* | 1 | -0.496 | 0.291 | 0.123 | | | -1.155 | 0.163 | 6 |
| ***M_dummy * WeeksGirdling::2*** | **2** | **-0.814** | **0.322** | **0.032** | * | | **-1.542** | **-0.086** | **14** |
| *M_dummy * WeeksGirdling::3* | 3 | -0.578 | 0.400 | 0.182 | | 0.562 | -1.483 | 0.327 | 20 |
| ***M_dummy * WeeksGirdling::4*** | **4** | **-0.795** | **0.348** | **0.048** | * | | **-1.582** | **-0.009** | **28** |
| *M_dummy * WeeksGirdling::5* | 5 | -0.564 | 0.567 | 0.346 | | | -1.846 | 0.719 | 41 |
| *M_dummy * WeeksGirdling::6* | 6 | -0.487 | 0.432 | 0.288 | | | -1.464 | 0.490 | 74 |

**E**

**Starch**

| Factors | Event | Estimate | Std. error | P-value | Sig. | Adj. R2 | 2.5% C.I. | 97.5% C.I. | Days girdling |
|---|---|---|---|---|---|---|---|---|---|
| *M_dummy * WeeksGirdling::0* | 0 | 0.000 | 0.000 | | | | 0.000 | 0.000 | -3 |
| *M_dummy * WeeksGirdling::1* | 1 | -1.883 | 2.425 | 0.457 | | | -7.369 | 3.603 | 6 |
| *M_dummy * WeeksGirdling::2* | 2 | 0.351 | 3.580 | 0.924 | | | -7.748 | 8.450 | 14 |
| *M_dummy * WeeksGirdling::3* | 3 | -1.390 | 2.876 | 0.640 | | 0.626 | -7.895 | 5.115 | 20 |
| *M_dummy * WeeksGirdling::4* | 4 | -0.274 | 3.711 | 0.943 | | | -8.669 | 8.122 | 28 |
| *M_dummy * WeeksGirdling::5* | 5 | -0.488 | 2.588 | 0.854 | | | -6.342 | 5.365 | 41 |
| *M_dummy * WeeksGirdling::6* | 6 | -0.705 | 1.157 | 0.558 | | | -3.323 | 1.914 | 74 |

**F**

**Total NSC**

| Factors | Event | Estimate | Std. error | P-value | Sig. | Adj. R2 | 2.5% C.I. | 97.5% C.I. | Days girdling |
|---|---|---|---|---|---|---|---|---|---|
| *M_dummy * WeeksGirdling::0* | 0 | 0.000 | 0.000 | | | | 0.000 | 0.000 | -3 |
| *M_dummy * WeeksGirdling::1* | 1 | -2.379 | 2.355 | 0.339 | | | -7.706 | 2.949 | 6 |
| *M_dummy * WeeksGirdling::2* | 2 | -0.463 | 3.533 | 0.899 | | | -8.456 | 7.530 | 14 |
| *M_dummy * WeeksGirdling::3* | 3 | -1.968 | 2.878 | 0.511 | | 0.668 | -8.477 | 4.541 | 20 |
| *M_dummy * WeeksGirdling::4* | 4 | -1.069 | 3.509 | 0.768 | | | -9.008 | 6.869 | 28 |
| *M_dummy * WeeksGirdling::5* | 5 | -1.052 | 2.743 | 0.710 | | | -7.257 | 5.153 | 41 |
| *M_dummy * WeeksGirdling::6* | 6 | -1.192 | 1.335 | 0.395 | | | -4.211 | 1.827 | 74 |

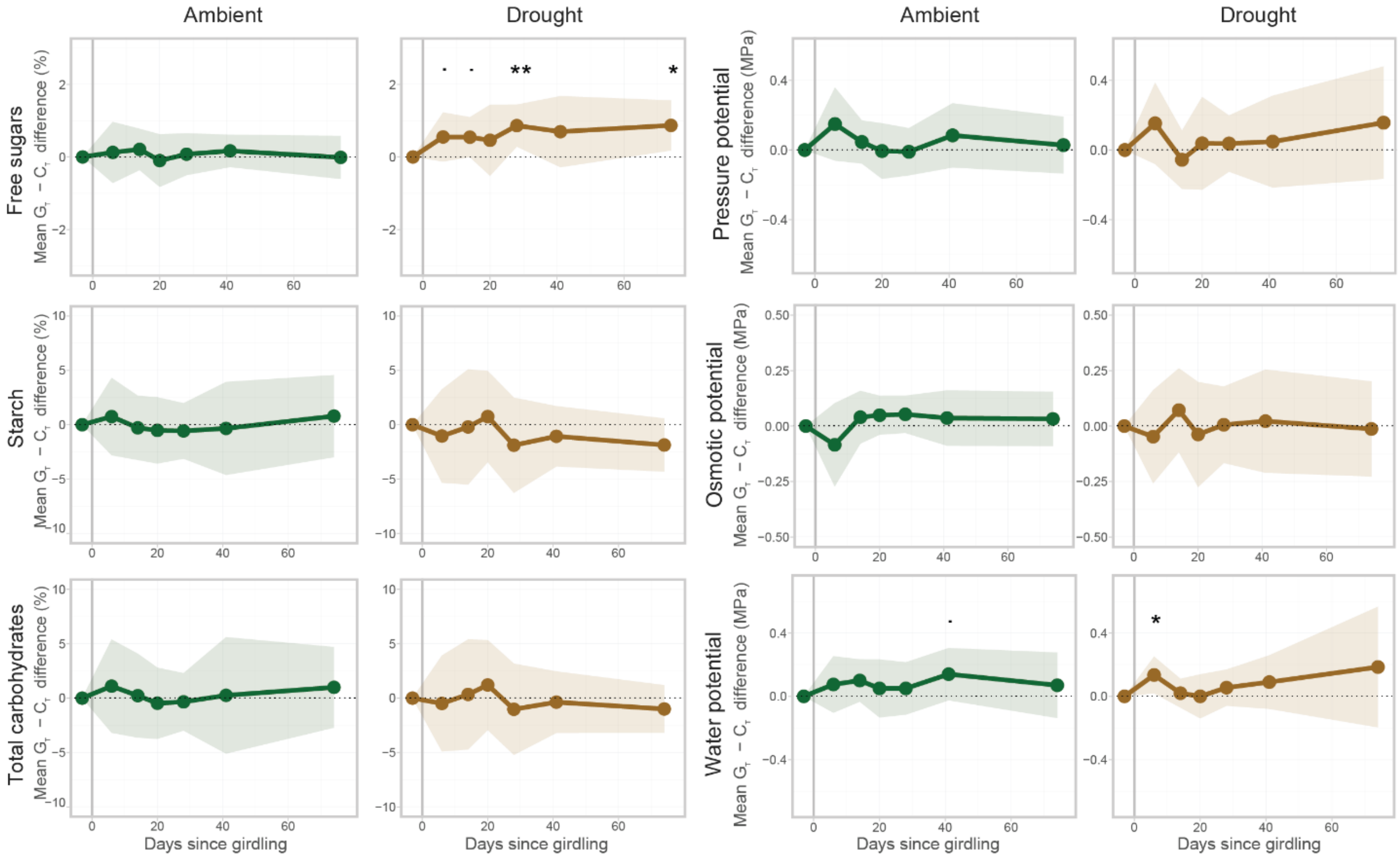


**Figure S6:** Panels show DiD estimates of treatment effects in non-structural carbohydrates (columns 1 and 2) and water relations (columns 3 and 4) under ambient (green) and drought (brown) conditions. Vertical line indicates the start of treatment. The DiD estimate is the difference in differences between trenched trees from girdled and control plots. Shaded areas represent 95% mean confidence intervals. Significance levels: · = $p < 0.1$, * = $p < 0.05$, ** = $p < 0.01$.

**Table S5:** Difference in differences models assessing changes in water relations variables in trenched trees due to girdling in ambient and drought plots. Bold highlights indicate significant effects. Significance levels: · = p < 0.1, * = p < 0.05.

**Ambient**

**A**

**Pressure potential**

| Factors | Event | Estimate | Std. error | P-value | Sig. | Adj. R2 | 2.5% C.I. | 97.5% C.I. | Days girdling |
|---|---|---|---|---|---|---|---|---|---|
| *M_dummy * WeeksGirdling::0* | 0 | 0.000 | 0.000 | | | | 0.000 | 0.000 | -3 |
| *M_dummy * WeeksGirdling::1* | 1 | 0.149 | 0.093 | 0.143 | | | -0.061 | 0.359 | 6 |
| *M_dummy * WeeksGirdling::2* | 2 | 0.046 | 0.055 | 0.422 | | | -0.078 | 0.170 | 14 |
| *M_dummy * WeeksGirdling::3* | 3 | -0.006 | 0.070 | 0.934 | | 0.649 | -0.165 | 0.153 | 20 |
| *M_dummy * WeeksGirdling::4* | 4 | -0.010 | 0.060 | 0.871 | | | -0.146 | 0.126 | 28 |
| *M_dummy * WeeksGirdling::5* | 5 | 0.084 | 0.081 | 0.329 | | | -0.100 | 0.268 | 41 |
| *M_dummy * WeeksGirdling::6* | 6 | 0.028 | 0.072 | 0.706 | | | -0.135 | 0.191 | 74 |

**B**

**Osmotic potential**

| Factors | Event | Estimate | Std. error | P-value | Sig. | Adj. R2 | 2.5% C.I. | 97.5% C.I. | Days girdling |
|---|---|---|---|---|---|---|---|---|---|
| *M_dummy * WeeksGirdling::0* | 0 | 0.000 | 0.000 | | | | 0.000 | 0.000 | -3 |
| *M_dummy * WeeksGirdling::1* | 1 | -0.085 | 0.083 | 0.334 | | | -0.272 | 0.103 | 6 |
| *M_dummy * WeeksGirdling::2* | 2 | 0.040 | 0.053 | 0.468 | | | -0.079 | 0.159 | 14 |
| *M_dummy * WeeksGirdling::3* | 3 | 0.049 | 0.039 | 0.239 | | 0.687 | -0.039 | 0.137 | 20 |
| *M_dummy * WeeksGirdling::4* | 4 | 0.053 | 0.037 | 0.188 | | | -0.031 | 0.137 | 28 |
| *M_dummy * WeeksGirdling::5* | 5 | 0.036 | 0.055 | 0.529 | | | -0.089 | 0.161 | 41 |
| *M_dummy * WeeksGirdling::6* | 6 | 0.032 | 0.054 | 0.568 | | | -0.090 | 0.155 | 74 |

**C**

**Water potential**

| Factors | Event | Estimate | Std. error | P-value | Sig. | Adj. R2 | 2.5% C.I. | 97.5% C.I. | Days girdling |
|---|---|---|---|---|---|---|---|---|---|
| *M_dummy * WeeksGirdling::0* | 0 | 0.000 | 0.000 | | | | 0.000 | 0.000 | -3 |
| *M_dummy * WeeksGirdling::1* | 1 | 0.075 | 0.079 | 0.366 | | | -0.103 | 0.253 | 6 |
| *M_dummy * WeeksGirdling::2* | 2 | 0.100 | 0.059 | 0.124 | | | -0.034 | 0.234 | 14 |
| *M_dummy * WeeksGirdling::3* | 3 | 0.050 | 0.081 | 0.552 | | 0.601 | -0.133 | 0.233 | 20 |
| *M_dummy * WeeksGirdling::4* | 4 | 0.050 | 0.073 | 0.511 | | | -0.115 | 0.215 | 28 |
| ***M_dummy * WeeksGirdling::5*** | **5** | **0.140** | **0.073** | **0.087** | **.** | | **-0.025** | **0.305** | **41** |
| *M_dummy * WeeksGirdling::6* | 6 | 0.070 | 0.091 | 0.463 | | | -0.137 | 0.277 | 74 |

### Table S5 (continued)

**Drought**

**D**

**Pressure potential**

| Factors | Event | Estimate | Std. error | P-value | Sig. | Adj. R2 | 2.5% C.I. | 97.5% C.I. | Days girdling |
|---|---|---|---|---|---|---|---|---|---|
| *M_dummy * WeeksGirdling::0* | 0 | 0.000 | 0.000 | | | | 0.000 | 0.000 | -3 |
| *M_dummy * WeeksGirdling::1* | 1 | 0.153 | 0.103 | 0.172 | | | -0.080 | 0.386 | 6 |
| *M_dummy * WeeksGirdling::2* | 2 | -0.056 | 0.074 | 0.468 | | | -0.223 | 0.111 | 14 |
| *M_dummy * WeeksGirdling::3* | 3 | 0.038 | 0.118 | 0.754 | | 0.616 | -0.228 | 0.304 | 20 |
| *M_dummy * WeeksGirdling::4* | 4 | 0.037 | 0.071 | 0.616 | | | -0.124 | 0.198 | 28 |
| *M_dummy * WeeksGirdling::5* | 5 | 0.048 | 0.117 | 0.690 | | | -0.216 | 0.312 | 41 |
| *M_dummy * WeeksGirdling::6* | 6 | 0.157 | 0.143 | 0.299 | | | -0.166 | 0.480 | 74 |

**E**

**Osmotic potential**

| Factors | Event | Estimate | Std. error | P-value | Sig. | Adj. R2 | 2.5% C.I. | 97.5% C.I. | Days girdling |
|---|---|---|---|---|---|---|---|---|---|
| *M_dummy * WeeksGirdling::0* | 0 | 0.000 | 0.000 | | | | 0.000 | 0.000 | -3 |
| *M_dummy * WeeksGirdling::1* | 1 | -0.048 | 0.093 | 0.618 | | | -0.258 | 0.162 | 6 |
| *M_dummy * WeeksGirdling::2* | 2 | 0.072 | 0.083 | 0.413 | | | -0.117 | 0.260 | 14 |
| *M_dummy * WeeksGirdling::3* | 3 | -0.038 | 0.105 | 0.725 | | 0.498 | -0.275 | 0.199 | 20 |
| *M_dummy * WeeksGirdling::4* | 4 | 0.006 | 0.076 | 0.941 | | | -0.166 | 0.178 | 28 |
| *M_dummy * WeeksGirdling::5* | 5 | 0.022 | 0.103 | 0.835 | | | -0.210 | 0.255 | 41 |
| *M_dummy * WeeksGirdling::6* | 6 | -0.013 | 0.095 | 0.894 | | | -0.227 | 0.201 | 74 |

**F**

**Water potential**

| Factors | Event | Estimate | Std. error | P-value | Sig. | Adj. R2 | 2.5% C.I. | 97.5% C.I. | Days girdling |
|---|---|---|---|---|---|---|---|---|---|
| *M_dummy * WeeksGirdling::0* | 0 | 0.000 | 0.000 | | | | 0.000 | 0.000 | -3 |
| ***M_dummy * WeeksGirdling::1*** | **1** | **0.135** | **0.051** | **0.027** | * | | **0.019** | **0.251** | **6** |
| *M_dummy * WeeksGirdling::2* | 2 | 0.020 | 0.040 | 0.631 | | | -0.071 | 0.111 | 14 |
| *M_dummy * WeeksGirdling::3* | 3 | 0.000 | 0.061 | 1.000 | | 0.694 | -0.139 | 0.139 | 20 |
| *M_dummy * WeeksGirdling::4* | 4 | 0.055 | 0.051 | 0.305 | | | -0.059 | 0.169 | 28 |
| *M_dummy * WeeksGirdling::5* | 5 | 0.090 | 0.075 | 0.259 | | | -0.079 | 0.259 | 41 |
| *M_dummy * WeeksGirdling::6* | 6 | 0.185 | 0.168 | 0.300 | | | -0.196 | 0.566 | 74 |

**Table S6:** Difference in differences models assessing changes in non-structural carbohydrate variables in trenched trees due to girdling in ambient and drought plots. Bold highlights indicate significant effects. Significance levels: · = p < 0.1, * = p < 0.05, ** = p < 0.01.

**Ambient**

**A**

**Soluble sugars**

| Factors | Event | Estimate | Std. error | P-value | Sig. | Adj. R2 | 2.5% C.I. | 97.5% C.I. | Days girdling |
|---|---|---|---|---|---|---|---|---|---|
| *M_dummy * WeeksGirdling::0* | 0 | 0.000 | 0.000 | | | | 0.000 | 0.000 | -3 |
| *M_dummy * WeeksGirdling::1* | 1 | 0.124 | 0.372 | 0.746 | | | -0.718 | 0.966 | 6 |
| *M_dummy * WeeksGirdling::2* | 2 | 0.207 | 0.252 | 0.432 | | | -0.362 | 0.776 | 14 |
| *M_dummy * WeeksGirdling::3* | 3 | -0.100 | 0.321 | 0.761 | | 0.899 | -0.826 | 0.625 | 20 |
| *M_dummy * WeeksGirdling::4* | 4 | 0.075 | 0.255 | 0.775 | | | -0.502 | 0.653 | 28 |
| *M_dummy * WeeksGirdling::5* | 5 | 0.167 | 0.197 | 0.420 | | | -0.279 | 0.613 | 41 |
| *M_dummy * WeeksGirdling::6* | 6 | -0.013 | 0.263 | 0.961 | | | -0.608 | 0.582 | 74 |

**B**

**Starch**

| Factors | Event | Estimate | Std. error | P-value | Sig. | Adj. R2 | 2.5% C.I. | 97.5% C.I. | Days girdling |
|---|---|---|---|---|---|---|---|---|---|
| *M_dummy * WeeksGirdling::0* | 0 | 0.000 | 0.000 | | | | 0.000 | 0.000 | -3 |
| *M_dummy * WeeksGirdling::1* | 1 | 0.749 | 1.571 | 0.645 | | | -2.806 | 4.303 | 6 |
| *M_dummy * WeeksGirdling::2* | 2 | -0.286 | 1.305 | 0.832 | | | -3.237 | 2.666 | 14 |
| *M_dummy * WeeksGirdling::3* | 3 | -0.520 | 1.346 | 0.708 | | 0.857 | -3.566 | 2.525 | 20 |
| *M_dummy * WeeksGirdling::4* | 4 | -0.570 | 1.127 | 0.625 | | | -3.120 | 1.980 | 28 |
| *M_dummy * WeeksGirdling::5* | 5 | -0.339 | 1.890 | 0.861 | | | -4.614 | 3.935 | 41 |
| *M_dummy * WeeksGirdling::6* | 6 | 0.793 | 1.668 | 0.646 | | | -2.981 | 4.567 | 74 |

**C**

**Total NSC**

| Factors | Event | Estimate | Std. error | P-value | Sig. | Adj. R2 | 2.5% C.I. | 97.5% C.I. | Days girdling |
|---|---|---|---|---|---|---|---|---|---|
| *M_dummy * WeeksGirdling::0* | 0 | 0.000 | 0.000 | | | | 0.000 | 0.000 | -3 |
| *M_dummy * WeeksGirdling::1* | 1 | 1.103 | 1.894 | 0.575 | | | -3.182 | 5.388 | 6 |
| *M_dummy * WeeksGirdling::2* | 2 | 0.228 | 1.704 | 0.897 | | | -3.628 | 4.084 | 14 |
| *M_dummy * WeeksGirdling::3* | 3 | -0.468 | 1.446 | 0.754 | | 0.870 | -3.739 | 2.804 | 20 |
| *M_dummy * WeeksGirdling::4* | 4 | -0.341 | 1.166 | 0.776 | | | -2.978 | 2.295 | 28 |
| *M_dummy * WeeksGirdling::5* | 5 | 0.256 | 2.366 | 0.916 | | | -5.095 | 5.608 | 41 |
| *M_dummy * WeeksGirdling::6* | 6 | 0.995 | 1.638 | 0.559 | | | -2.710 | 4.700 | 74 |

### Table S6 (continued)

**Drought**

**D**

**Soluble sugars**

| Factors | Event | Estimate | Std. error | P-value | Sig. | Adj. R2 | 2.5% C.I. | 97.5% C.I. | Days girdling |
|---|---|---|---|---|---|---|---|---|---|
| *M_dummy * WeeksGirdling::0* | 0 | 0.000 | 0.000 | | | | 0.000 | 0.000 | -3 |
| ***M_dummy * WeeksGirdling::1*** | **1** | **0.551** | **0.299** | **0.099** | **.** | | **-0.125** | **1.227** | **6** |
| ***M_dummy * WeeksGirdling::2*** | **2** | **0.545** | **0.244** | **0.052** | **.** | | **-0.007** | **1.096** | **14** |
| *M_dummy * WeeksGirdling::3* | 3 | 0.459 | 0.433 | 0.316 | | 0.590 | -0.520 | 1.438 | 20 |
| ***M_dummy * WeeksGirdling::4*** | **4** | **0.863** | **0.256** | **0.008** | ** | | **0.285** | **1.442** | **28** |
| *M_dummy * WeeksGirdling::5* | 5 | 0.697 | 0.433 | 0.142 | | | -0.284 | 1.678 | 41 |
| ***M_dummy * WeeksGirdling::6*** | **6** | **0.869** | **0.305** | **0.019** | * | | **0.179** | **1.559** | **74** |

**E**

**Starch**

| Factors | Event | Estimate | Std. error | P-value | Sig. | Adj. R2 | 2.5% C.I. | 97.5% C.I. | Days girdling |
|---|---|---|---|---|---|---|---|---|---|
| *M_dummy * WeeksGirdling::0* | 0 | 0.000 | 0.000 | | | | 0.000 | 0.000 | -3 |
| *M_dummy * WeeksGirdling::1* | 1 | -1.039 | 1.897 | 0.597 | | | -5.329 | 3.252 | 6 |
| *M_dummy * WeeksGirdling::2* | 2 | -0.203 | 2.338 | 0.933 | | | -5.493 | 5.086 | 14 |
| *M_dummy * WeeksGirdling::3* | 3 | 0.752 | 1.851 | 0.694 | | 0.869 | -3.436 | 4.939 | 20 |
| *M_dummy * WeeksGirdling::4* | 4 | -1.884 | 1.935 | 0.356 | | | -6.260 | 2.492 | 28 |
| *M_dummy * WeeksGirdling::5* | 5 | -1.064 | 1.225 | 0.408 | | | -3.835 | 1.707 | 41 |
| *M_dummy * WeeksGirdling::6* | 6 | -1.864 | 1.081 | 0.119 | | | -4.310 | 0.583 | 74 |

**F**

**Total NSC**

| Factors | Event | Estimate | Std. error | P-value | Sig. | Adj. R2 | 2.5% C.I. | 97.5% C.I. | Days girdling |
|---|---|---|---|---|---|---|---|---|---|
| *M_dummy * WeeksGirdling::0* | 0 | 0.000 | 0.000 | | | | 0.000 | 0.000 | -3 |
| *M_dummy * WeeksGirdling::1* | 1 | -0.488 | 1.947 | 0.808 | | | -4.893 | 3.916 | 6 |
| *M_dummy * WeeksGirdling::2* | 2 | 0.342 | 2.236 | 0.882 | | | -4.717 | 5.400 | 14 |
| *M_dummy * WeeksGirdling::3* | 3 | 1.211 | 1.827 | 0.524 | | 0.887 | -2.923 | 5.344 | 20 |
| *M_dummy * WeeksGirdling::4* | 4 | -1.021 | 1.854 | 0.595 | | | -5.215 | 3.173 | 28 |
| *M_dummy * WeeksGirdling::5* | 5 | -0.367 | 1.249 | 0.776 | | | -3.191 | 2.458 | 41 |
| *M_dummy * WeeksGirdling::6* | 6 | -0.995 | 0.968 | 0.331 | | | -3.186 | 1.196 | 74 |

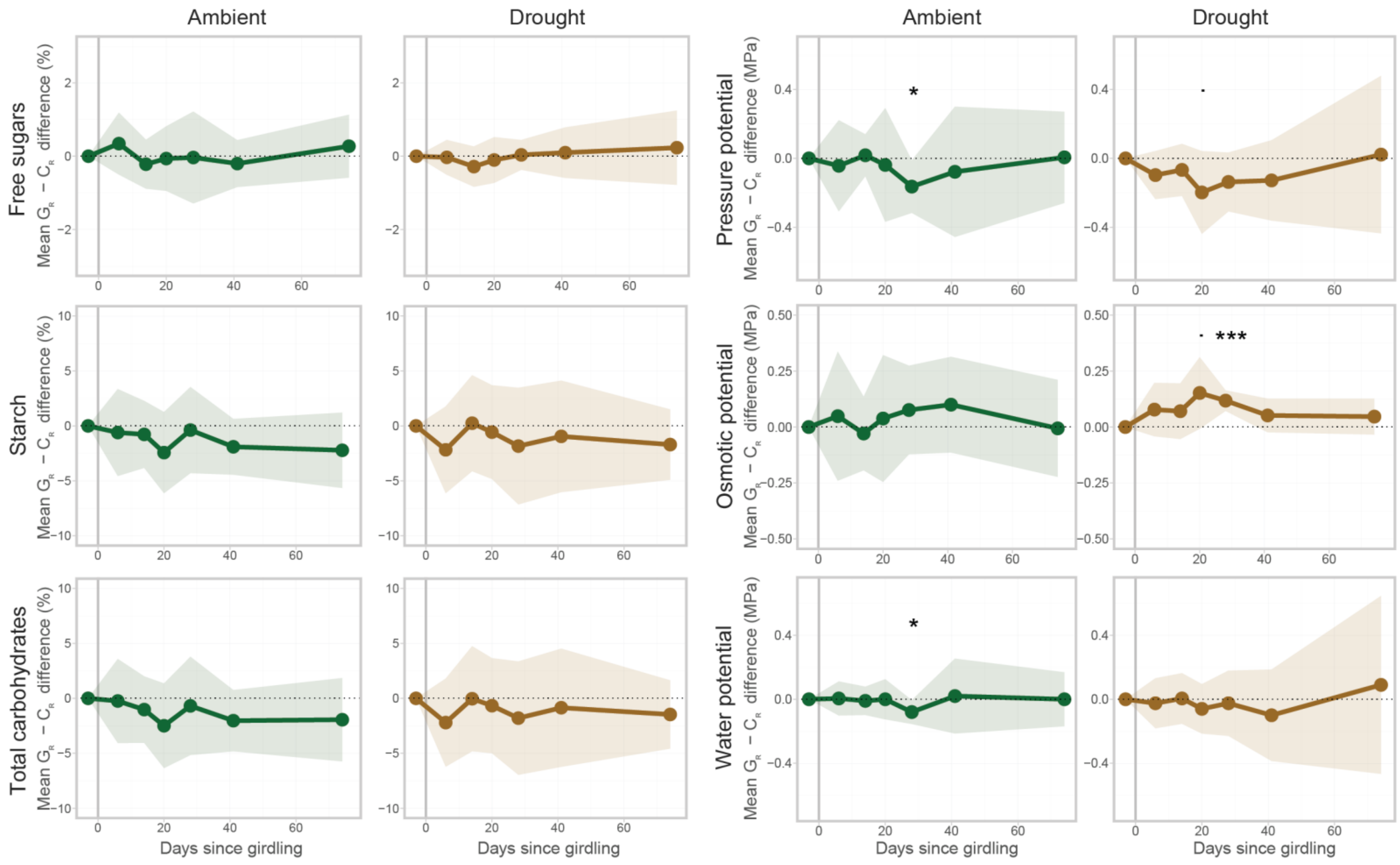


**Figure S7:** Panels show DiD estimates of treatment effects in non-structural carbohydrates (columns 1 and 2) and water relations (columns 3 and 4) under ambient (green) and drought (brown) conditions. Vertical line indicates the start of treatment. The DiD estimate is the difference in differences between response trees from girdled and control plots. Shaded areas represent 95% mean confidence intervals. Significance levels: · = $p < 0.1$, * = $p < 0.05$, *** = $p < 0.001$.

**Table S7:** Difference in differences models assessing changes in water relations variables in response trees due to girdling in ambient and drought plots. Bold highlights indicate significant effects. Significance levels: · = p < 0.1, * = p < 0.05, *** = p < 0.001.

**Ambient**

**A**

**Pressure potential**

| Factors | Event | Estimate | Std. error | P-value | Sig. | Adj. R2 | 2.5% C.I. | 97.5% C.I. | Days girdling |
|---|---|---|---|---|---|---|---|---|---|
| *M_dummy * WeeksGirdling::0* | 0 | 0.000 | 0.000 | | | | 0.000 | 0.000 | -3 |
| *M_dummy * WeeksGirdling::1* | 1 | -0.043 | 0.117 | 0.722 | | | -0.308 | 0.222 | 6 |
| *M_dummy * WeeksGirdling::2* | 2 | 0.018 | 0.054 | 0.746 | | | -0.104 | 0.140 | 14 |
| *M_dummy * WeeksGirdling::3* | 3 | -0.038 | 0.146 | 0.800 | | 0.639 | -0.368 | 0.292 | 20 |
| ***M_dummy * WeeksGirdling::4*** | **4** | **-0.164** | **0.067** | **0.038** | * | | **-0.317** | **-0.011** | **28** |
| *M_dummy * WeeksGirdling::5* | 5 | -0.078 | 0.167 | 0.652 | | | -0.456 | 0.300 | 41 |
| *M_dummy * WeeksGirdling::6* | 6 | 0.006 | 0.117 | 0.960 | | | -0.259 | 0.271 | 74 |

**B**

**Osmotic potential**

| Factors | Event | Estimate | Std. error | P-value | Sig. | Adj. R2 | 2.5% C.I. | 97.5% C.I. | Days girdling |
|---|---|---|---|---|---|---|---|---|---|
| *M_dummy * WeeksGirdling::0* | 0 | 0.000 | 0.000 | | | | 0.000 | 0.000 | -3 |
| *M_dummy * WeeksGirdling::1* | 1 | 0.048 | 0.127 | 0.712 | | | -0.240 | 0.337 | 6 |
| *M_dummy * WeeksGirdling::2* | 2 | -0.029 | 0.073 | 0.699 | | | -0.193 | 0.135 | 14 |
| *M_dummy * WeeksGirdling::3* | 3 | 0.038 | 0.125 | 0.768 | | 0.612 | -0.245 | 0.321 | 20 |
| *M_dummy * WeeksGirdling::4* | 4 | 0.076 | 0.088 | 0.407 | | | -0.122 | 0.274 | 28 |
| *M_dummy * WeeksGirdling::5* | 5 | 0.100 | 0.095 | 0.318 | | | -0.114 | 0.314 | 41 |
| *M_dummy * WeeksGirdling::6* | 6 | -0.006 | 0.096 | 0.951 | | | -0.223 | 0.211 | 74 |

**C**

**Water potential**

| Factors | Event | Estimate | Std. error | P-value | Sig. | Adj. R2 | 2.5% C.I. | 97.5% C.I. | Days girdling |
|---|---|---|---|---|---|---|---|---|---|
| *M_dummy * WeeksGirdling::0* | 0 | 0.000 | 0.000 | | | | 0.000 | 0.000 | -3 |
| *M_dummy * WeeksGirdling::1* | 1 | 0.005 | 0.047 | 0.918 | | | -0.102 | 0.112 | 6 |
| *M_dummy * WeeksGirdling::2* | 2 | -0.010 | 0.039 | 0.804 | | | -0.099 | 0.079 | 14 |
| *M_dummy * WeeksGirdling::3* | 3 | 0.000 | 0.055 | 1.000 | | 0.654 | -0.126 | 0.126 | 20 |
| ***M_dummy * WeeksGirdling::4*** | **4** | **-0.080** | **0.033** | **0.038** | * | | **-0.155** | **-0.005** | **28** |
| *M_dummy * WeeksGirdling::5* | 5 | 0.020 | 0.104 | 0.851 | | | -0.214 | 0.254 | 41 |
| *M_dummy * WeeksGirdling::6* | 6 | 0.000 | 0.075 | 1.000 | | | -0.169 | 0.169 | 74 |

### Table S7 (continued)

**Drought**

**D**

**Pressure potential**

| Factors | Event | Estimate | Std. error | P-value | Sig. | Adj. R2 | 2.5% C.I. | 97.5% C.I. | Days girdling |
|---|---|---|---|---|---|---|---|---|---|
| M_dummy * WeeksGirdling::0 | 0 | 0.000 | 0.000 | | | | 0.000 | 0.000 | -3 |
| M_dummy * WeeksGirdling::1 | 1 | -0.097 | 0.062 | 0.151 | | | -0.237 | 0.043 | 6 |
| M_dummy * WeeksGirdling::2 | 2 | -0.067 | 0.067 | 0.343 | | | -0.218 | 0.084 | 14 |
| **M_dummy * WeeksGirdling::3** | **3** | **-0.198** | **0.106** | **0.095** | **.** | 0.716 | **-0.438** | **0.042** | **20** |
| M_dummy * WeeksGirdling::4 | 4 | -0.137 | 0.076 | 0.104 | | | -0.308 | 0.034 | 28 |
| M_dummy * WeeksGirdling::5 | 5 | -0.128 | 0.103 | 0.246 | | | -0.362 | 0.106 | 41 |
| M_dummy * WeeksGirdling::6 | 6 | 0.022 | 0.202 | 0.916 | | | -0.436 | 0.480 | 74 |

**E**

**Osmotic potential**

| Factors | Event | Estimate | Std. error | P-value | Sig. | Adj. R2 | 2.5% C.I. | 97.5% C.I. | Days girdling |
|---|---|---|---|---|---|---|---|---|---|
| M_dummy * WeeksGirdling::0 | 0 | 0.000 | 0.000 | | | | 0.000 | 0.000 | -3 |
| M_dummy * WeeksGirdling::1 | 1 | 0.078 | 0.053 | 0.174 | | | -0.042 | 0.197 | 6 |
| M_dummy * WeeksGirdling::2 | 2 | 0.071 | 0.055 | 0.230 | | | -0.054 | 0.195 | 14 |
| **M_dummy * WeeksGirdling::3** | **3** | **0.152** | **0.071** | **0.060** | **.** | 0.744 | **-0.008** | **0.312** | **20** |
| **M_dummy * WeeksGirdling::4** | **4** | **0.118** | **0.020** | **0.000** | ******* | | **0.072** | **0.164** | **28** |
| M_dummy * WeeksGirdling::5 | 5 | 0.052 | 0.033 | 0.154 | | | -0.023 | 0.127 | 41 |
| M_dummy * WeeksGirdling::6 | 6 | 0.047 | 0.035 | 0.222 | | | -0.034 | 0.127 | 74 |

**F**

**Water potential**

| Factors | Event | Estimate | Std. error | P-value | Sig. | Adj. R2 | 2.5% C.I. | 97.5% C.I. | Days girdling |
|---|---|---|---|---|---|---|---|---|---|
| M_dummy * WeeksGirdling::0 | 0 | 0.000 | 0.000 | | | | 0.000 | 0.000 | -3 |
| M_dummy * WeeksGirdling::1 | 1 | -0.025 | 0.069 | 0.726 | | | -0.182 | 0.132 | 6 |
| M_dummy * WeeksGirdling::2 | 2 | 0.005 | 0.070 | 0.945 | | | -0.154 | 0.164 | 14 |
| M_dummy * WeeksGirdling::3 | 3 | -0.060 | 0.068 | 0.403 | | 0.743 | -0.215 | 0.095 | 20 |
| M_dummy * WeeksGirdling::4 | 4 | -0.025 | 0.090 | 0.788 | | | -0.229 | 0.179 | 28 |
| M_dummy * WeeksGirdling::5 | 5 | -0.100 | 0.127 | 0.450 | | | -0.386 | 0.186 | 41 |
| M_dummy * WeeksGirdling::6 | 6 | 0.090 | 0.246 | 0.723 | | | -0.467 | 0.647 | 74 |

**Table S8:** Difference in differences models assessing changes in non-structural carbohydrate variables in response trees due to girdling in ambient and drought plots. Bold highlights indicate significant effects.

**Ambient**

**A**

**Soluble sugars**

| **Factors** | **Event** | **Estimate** | **Std. error** | **P-value** | **Sig.** | **Adj. R2** | **2.5% C.I.** | **97.5% C.I.** | **Days girdling** |
|---|---|---|---|---|---|---|---|---|---|
| *M_dummy * WeeksGirdling::0* | 0 | 0.000 | 0.000 | | | | 0.000 | 0.000 | -3 |
| *M_dummy * WeeksGirdling::1* | 1 | 0.344 | 0.374 | 0.382 | | | -0.502 | 1.189 | 6 |
| *M_dummy * WeeksGirdling::2* | 2 | -0.218 | 0.295 | 0.479 | | | -0.886 | 0.450 | 14 |
| *M_dummy * WeeksGirdling::3* | 3 | -0.065 | 0.389 | 0.872 | | 0.804 | -0.946 | 0.816 | 20 |
| *M_dummy * WeeksGirdling::4* | 4 | -0.036 | 0.554 | 0.950 | | | -1.289 | 1.218 | 28 |
| *M_dummy * WeeksGirdling::5* | 5 | -0.201 | 0.283 | 0.495 | | | -0.841 | 0.439 | 41 |
| *M_dummy * WeeksGirdling::6* | 6 | 0.273 | 0.380 | 0.491 | | | -0.586 | 1.131 | 74 |

**B**

**Starch**

| **Factors** | **Event** | **Estimate** | **Std. error** | **P-value** | **Sig.** | **Adj. R2** | **2.5% C.I.** | **97.5% C.I.** | **Days girdling** |
|---|---|---|---|---|---|---|---|---|---|
| *M_dummy * WeeksGirdling::0* | 0 | 0.000 | 0.000 | | | | 0.000 | 0.000 | -3 |
| *M_dummy * WeeksGirdling::1* | 1 | -0.602 | 1.748 | 0.738 | | | -4.556 | 3.351 | 6 |
| *M_dummy * WeeksGirdling::2* | 2 | -0.773 | 1.347 | 0.580 | | | -3.821 | 2.275 | 14 |
| *M_dummy * WeeksGirdling::3* | 3 | -2.430 | 1.632 | 0.171 | | 0.876 | -6.122 | 1.262 | 20 |
| *M_dummy * WeeksGirdling::4* | 4 | -0.378 | 1.732 | 0.832 | | | -4.295 | 3.539 | 28 |
| *M_dummy * WeeksGirdling::5* | 5 | -1.900 | 1.122 | 0.125 | | | -4.438 | 0.638 | 41 |
| *M_dummy * WeeksGirdling::6* | 6 | -2.218 | 1.516 | 0.177 | | | -5.647 | 1.211 | 74 |

**C**

**Total NSC**

| **Factors** | **Event** | **Estimate** | **Std. error** | **P-value** | **Sig.** | **Adj. R2** | **2.5% C.I.** | **97.5% C.I.** | **Days girdling** |
|---|---|---|---|---|---|---|---|---|---|
| *M_dummy * WeeksGirdling::0* | 0 | 0.000 | 0.000 | | | | 0.000 | 0.000 | -3 |
| *M_dummy * WeeksGirdling::1* | 1 | -0.242 | 1.690 | 0.889 | | | -4.064 | 3.581 | 6 |
| *M_dummy * WeeksGirdling::2* | 2 | -1.025 | 1.337 | 0.463 | | | -4.050 | 1.999 | 14 |
| *M_dummy * WeeksGirdling::3* | 3 | -2.495 | 1.702 | 0.177 | | 0.895 | -6.346 | 1.357 | 20 |
| *M_dummy * WeeksGirdling::4* | 4 | -0.687 | 1.975 | 0.736 | | | -5.155 | 3.782 | 28 |
| *M_dummy * WeeksGirdling::5* | 5 | -2.033 | 1.232 | 0.133 | | | -4.819 | 0.754 | 41 |
| *M_dummy * WeeksGirdling::6* | 6 | -1.945 | 1.676 | 0.276 | | | -5.736 | 1.846 | 74 |

### Table S8 (continued)

**Drought**

**D**

**Soluble sugars**

| Factors | Event | Estimate | Std. error | P-value | Sig. | Adj. R2 | 2.5% C.I. | 97.5% C.I. | Days girdling |
|---|---|---|---|---|---|---|---|---|---|
| *M_dummy * WeeksGirdling::0* | 0 | 0.000 | 0.000 | | | | 0.000 | 0.000 | -3 |
| *M_dummy * WeeksGirdling::1* | 1 | -0.032 | 0.210 | 0.884 | | | -0.506 | 0.443 | 6 |
| *M_dummy * WeeksGirdling::2* | 2 | -0.284 | 0.243 | 0.273 | | | -0.834 | 0.266 | 14 |
| *M_dummy * WeeksGirdling::3* | 3 | -0.103 | 0.276 | 0.717 | | 0.814 | -0.728 | 0.521 | 20 |
| *M_dummy * WeeksGirdling::4* | 4 | 0.036 | 0.180 | 0.844 | | | -0.371 | 0.444 | 28 |
| *M_dummy * WeeksGirdling::5* | 5 | 0.097 | 0.305 | 0.758 | | | -0.593 | 0.787 | 41 |
| *M_dummy * WeeksGirdling::6* | 6 | 0.232 | 0.448 | 0.618 | | | -0.782 | 1.246 | 74 |

**E**

**Starch**

| Factors | Event | Estimate | Std. error | P-value | Sig. | Adj. R2 | 2.5% C.I. | 97.5% C.I. | Days girdling |
|---|---|---|---|---|---|---|---|---|---|
| *M_dummy * WeeksGirdling::0* | 0 | 0.000 | 0.000 | | | | 0.000 | 0.000 | -3 |
| *M_dummy * WeeksGirdling::1* | 1 | -2.176 | 1.745 | 0.244 | | | -6.124 | 1.773 | 6 |
| *M_dummy * WeeksGirdling::2* | 2 | 0.247 | 1.930 | 0.901 | | | -4.119 | 4.613 | 14 |
| *M_dummy * WeeksGirdling::3* | 3 | -0.564 | 1.886 | 0.772 | | 0.867 | -4.831 | 3.702 | 20 |
| *M_dummy * WeeksGirdling::4* | 4 | -1.836 | 2.346 | 0.454 | | | -7.143 | 3.472 | 28 |
| *M_dummy * WeeksGirdling::5* | 5 | -0.951 | 2.243 | 0.681 | | | -6.025 | 4.122 | 41 |
| *M_dummy * WeeksGirdling::6* | 6 | -1.699 | 1.416 | 0.261 | | | -4.903 | 1.505 | 74 |

**F**

**Total NSC**

| Factors | Event | Estimate | Std. error | P-value | Sig. | Adj. R2 | 2.5% C.I. | 97.5% C.I. | Days girdling |
|---|---|---|---|---|---|---|---|---|---|
| *M_dummy * WeeksGirdling::0* | 0 | 0.000 | 0.000 | | | | 0.000 | 0.000 | -3 |
| *M_dummy * WeeksGirdling::1* | 1 | -2.207 | 1.773 | 0.245 | | | -6.217 | 1.803 | 6 |
| *M_dummy * WeeksGirdling::2* | 2 | -0.037 | 2.114 | 0.987 | | | -4.819 | 4.746 | 14 |
| *M_dummy * WeeksGirdling::3* | 3 | -0.668 | 1.912 | 0.735 | | 0.879 | -4.992 | 3.657 | 20 |
| *M_dummy * WeeksGirdling::4* | 4 | -1.799 | 2.280 | 0.450 | | | -6.957 | 3.358 | 28 |
| *M_dummy * WeeksGirdling::5* | 5 | -0.855 | 2.376 | 0.727 | | | -6.230 | 4.521 | 41 |
| *M_dummy * WeeksGirdling::6* | 6 | -1.468 | 1.373 | 0.313 | | | -4.574 | 1.639 | 74 |

### Appendix S1

#### *Root grafting*

The goal of this experiment was to look explicitly for root grafts. We selected four sites with a treatment *Pinus ponderosa* tree between 5 and 13 cm of diameter at breast height near neighboring trees of the same species. There was a distance of two meters or less between the treatment tree and at least one neighboring tree.

*Methods*

On August 22nd, we cut and applied the 0.1% acid fuchsin dye treatment to all four treatment trees[1]. We chose the cut height of our treatment tree to best fit the rubber collar connecting our pvc reservoir to the trunk. Parafilm and silicone tape were used to fit the rubber collar tightly against the cut and peeled xylem column, followed by a coat of pruners paint around the interface with the tree (Fig 3, a, b). Any branches below the cut site were removed and the cuts were sealed with pruners paint to prevent leakage. Then, the reservoir was filled with dye. The next day, additional dye solution was added to the reservoir. Ahead of sampling, we kept the reservoir filled with water to maintain the backward flow of water through the treatment tree.

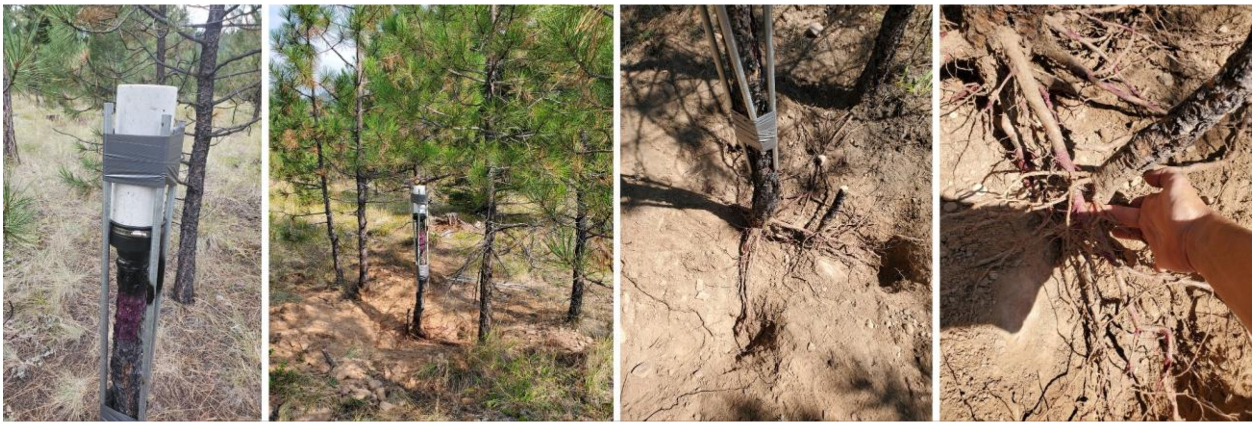

A B C D

***Figure 3.*** *A) Reservoir set-up on a tree for method testing. B) Coarse roots were excavated to look for grafts between the treatment and four neighboring trees. C) The excavated bole of the test tree was infused with dye which did not enter the soil. D) The bole of the test tree and the nearest neighbor sapling had intertwined coarse roots without evidence of grafting or dye transfer.*

On August 24th, we began root excavation at three of the four sites. The fourth site was left intact to allow more time for the dye to travel, as it had been deployed last the previous day. Additional water was added to the fourth reservoir the following day as well. Coarse roots were located at the bole of the treatment tree, and any roots in the 90-degree section of the treatment tree facing an adjacent tree were excavated and traced outward until they plunged to a depth below 40 cm (Fig. 4).

On October 2nd, 40 days after excavation, we used an increment borer to core the adjacent trees at breast height in all four replicate sites to inspect for dye traces.

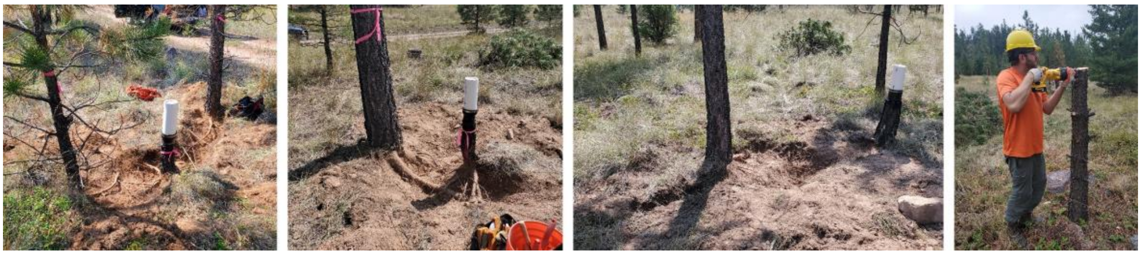

A B C D

***Figure 4.*** *Excavation of A) treatment tree #1, B) treatment tree #2, and C) treatment tree #3. D) Preparatory steps for treatment tree #4 involved the installation of the dye reservoir slightly above breast height to accommodate the size of the rubber collar best. We allowed additional time for the dye to travel for tree #4 to account for the added length.*

*Results*

No root grafts were found in any of the treatment sites at 40 cm or above. Some crossings of root systems were identified, but any overlapping coarse roots had not fused. We found no evidence of dye in any neighboring tree roots (Fig. 5). Follow up cores of closest adjacent trees taken with an increment borer 40 days after excavation showed no indication of acid fuchsin uptake in any of the four experiment sites.

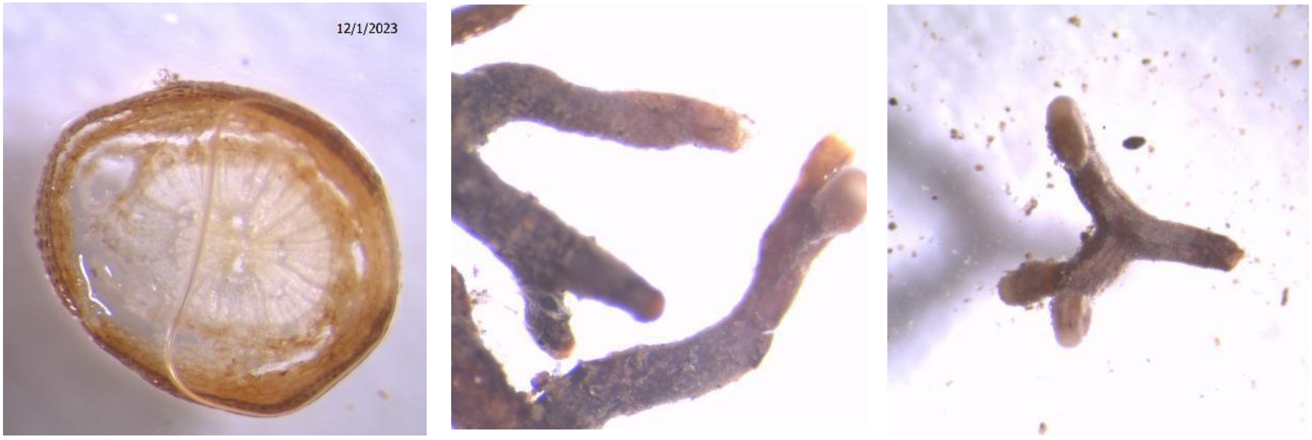


A B C

***Figure 5.*** *No dye observed in A) neighboring fine roots or B & C) root tips.*
